# PRDM1 Drives Chemoradiotherapy-associated Enrichment of Adaptive NK Cells in Cervical Cancer

**DOI:** 10.64898/2026.01.28.702427

**Authors:** Meng Wan, Tangwu Zhong, Wenyang Shi, Jianyu Shen, Wei Zhang, Yizhe Sun

**Affiliations:** Department of Radiation Oncology, Beijing Obstetrics and Gynecology Hospital, Capital Medical University; Beijing Maternal and Child Health Care Hospital, Beijing, China; School of Basic Medicine, Jiamusi University, Jiamusi, Heilongjiang, China; Department of Laboratory Medicine, Division of Pathology, Karolinska Institutet, Stockholm, Sweden; Science for Life Laboratory, Department of Oncology-Pathology, Karolinska Institutet, Stockholm, Sweden; Department of Clinical Science, Karolinska Institutet, Stockholm, Sweden

**Keywords:** Chemotherapy, Cervical cancer, adaptive NK cells, *In silico* perturbation, PRDM1

## Abstract

Chemoradiotherapy (CRT) induces not only direct tumor cell death but also extensive remodeling of the tumor immune microenvironment. Adaptive natural killer (aNK) cells, initially characterized in chronic viral infection, are increasingly recognized as functionally relevant immune populations in solid tumors, where they may contribute to antitumor immunity and immune memory. However, how aNK cells dynamically respond to chemoradiotherapy and the regulatory mechanisms underlying their activation in solid tumors remain poorly defined.

To address this, we analyzed single-cell RNA sequencing data from cervical cancer patients collected before CRT, after the first CRT fraction, and after the second fraction. Single-cell profiling revealed a significant enrichment of aNK cells following CRT. Differential gene expression and pathway enrichment analyses demonstrated that CRT-associated aNK cells exhibit enhanced virus-defending programs with increased cytotoxicity. Among the differentially expressed genes, the transcription factor PRDM1 was consistently and robustly upregulated in aNK cells after both the first and second rounds of CRT.

To investigate the functional role of PRDM1, we applied *in silico* perturbation analyses using scTenifoldKnk and CellOracle. Virtual knockout of PRDM1 resulted in a marked attenuation of effector programs and disruption of metabolic networks in aNK cells. Moreover, PRDM1 perturbation altered inferred cellular trajectories, reversing the progression toward the aNK cell state, suggesting a requirement for PRDM1 in maintaining aNK identity and functional maturation within the CRT-conditioned tumor microenvironment.

Together, these findings identify PRDM1 as a key regulatory factor associated with the enrichment, functional activation, and trajectory stabilization of aNK cells following CRT in cervical cancer, providing insight into innate immune remodeling during CRT and highlighting PRDM1 as a potential target for enhancing radiotherapy-induced antitumor immunity.

## Introduction

Chemoradiotherapy (CRT) remains a cornerstone treatment for patients with locally advanced cervical cancer and constitutes the standard of care for definitive management of this disease^[1, 2]^. Beyond its direct cytotoxic effects mediated by DNA damage, cell cycle arrest, and apoptosis, accumulating evidence indicates that CRT profoundly remodels the tumor immune microenvironment^[3]^. Both radiotherapy and chemotherapy can induce immunogenic cell death characterized by the release of damage-associated molecular patterns (DAMPs), including HMGB1, ATP, and surface calreticulin exposure, which promote dendritic cell activation and antigen uptake, thereby enhancing tumor antigen presentation and priming of antitumor T cell responses^[4, 5]^. In parallel, radiation increases the expression of MHC class I molecules, death receptors, and stress-induced ligands on tumor cells, augmenting their susceptibility to immune-mediated killing by cytotoxic lymphocytes, including CD8^+^ T cells and natural killer (NK) cells^[6, 7]^. Chemotherapeutic agents further synergize with radiotherapy by modulating immune checkpoint pathways, depleting immunosuppressive cell populations such as regulatory T cells and myeloid-derived suppressor cells, and promoting a pro-inflammatory cytokine milieu within the tumor microenvironment^[8-10]^. Collectively, these therapy-induced immune alterations drive recruitment, activation, and functional reprogramming of diverse immune cell subsets, ultimately shaping therapeutic efficacy and long-term clinical outcomes^[11, 12]^.

NK cells are key innate lymphocytes involved in tumor surveillance and antiviral defense^[13]^. In the setting of chemoradiotherapy, NK cell states are shaped by therapy-induced inflammatory cues and stress signaling in the tumor microenvironment: radiation can trigger cytosolic DNA sensing and cGAS–STING–dependent type I interferon programs that promote immune activation and can potentiate NK-cell activity, while also inducing chemokines that support immune-cell recruitment^[14, 15]^. At the tumor-cell level, irradiation can increase expression of immune-relevant surface molecules, including stress-induced ligands such as NKG2D ligands and death receptors, thereby enhancing susceptibility to NK-mediated cytotoxicity, although opposing effects (e.g., altered checkpoint/ligand balance) have also been reported depending on dose and context^[16]^. Chemotherapy can further modulate NK responses by inducing cellular stress pathways in malignant cells and remodeling suppressive or inflammatory components of the tumor milieu, creating conditions that alter NK activation thresholds and effector outputs^[17]^. In cervical cancer specifically, single-cell studies of radiochemotherapy/concurrent CRT have demonstrated marked immune-ecosystem remodeling after treatment, underscoring that NK-cell activation states and regulatory programs are dynamically rewired during therapy^[18]^. Commonly, transcriptional regulation is a key determinant of NK cell differentiation, functional maturation, and context-dependent activation within tumors. These therapy-conditioned signals are integrated by core NK transcriptional circuits, with T-bet (TBX21) and Eomes (EOMES) functioning as key maturation checkpoints, and by downstream regulators such as PRDM1 (Blimp-1), which controls NK cell maturation and effector cytokine programs^[19-21]^.In summary, these observations support a model in which CRT reshapes the cervical tumor microenvironment to dynamically reprogram NK cells through inflammatory and stress-associated pathways, altering both tumor susceptibility to NK killing and NK activation thresholds.

Beyond conventional NK cells, a distinct subset termed adaptive or memory-like NK (aNK) cells was first described in the context of chronic viral infection, most prominently human cytomegalovirus (HCMV), where virus-driven selection and expansion generate a highly differentiated NK population with a characteristic receptor and signaling profile ^[22-24]^. These cells are supported by stable epigenetic remodeling and long-term persistence, and can display clonal-like expansion with augmented effector potential, particularly in antibody-dependent cellular cytotoxicity and recall-like responsiveness ^[25-27]^ . Importantly, the relevance of aNK cells is increasingly recognized beyond infection. Multiple studies and reviews have highlighted their presence in cancer and their potential contribution to antitumor immunity, including enhanced cytotoxic competence and therapeutic promise in solid-tumor settings, especially in contexts where antibody-dependent mechanisms are engaged^[28, 29]^. Nevertheless, whether and how aNK cells dynamically respond to chemoradiotherapy in human cancers has not been systematically explored.

In this study, we analyzed single-cell RNA sequencing data from cervical cancer samples collected before CRT and at sequential time points during treatment to characterize therapy-induced immune remodeling. We observed an enrichment of adaptive NK cells after CRT together with activation of cytotoxic and antiviral transcriptional programs, and identified PRDM1 as a prominently induced transcriptional regulator in this population. To probe the functional relevance of PRDM1, we applied complementary *in silico* perturbation approaches to infer regulatory and trajectory-level consequences of PRDM1 disruption. Together, these analyses support a model in which PRDM1 contributes to CRT-associated adaptive NK cell responses and provide a framework for understanding transcriptional control of therapy-driven NK-cell adaptation in cervical cancer.

## Results

### CRT is associated with enrichment of aNK cells and induction of PRDM1 in aNK cells

We analyzed publicly available single-cell RNA-sequencing (scRNA-seq) data from cervical cancer patients in the GEO dataset GSE297041^[30]^, including samples collected before CRT (preRT), after the first CRT fraction (onRT1), and after the second fraction (onRT2). From the global immune landscape, NK cells were first identified and extracted based on NK features and visualized on a UMAP embedding, showing a distinct NK compartment separated from other immune populations **(Fig. 1A)**. Next, to annotate all NK cell clusters **(Fig.1B)**, we applied **scType**^[31]^ to identify the cluster most consistent with **aNK features**, using marker genes compiled from two sources: bulk RNA-seq of late-mature aNK versus early-mature cNK cells from human peripheral blood^[32, 33]^ and transcriptomic profiles of ovarian tumor– infiltrating NK cells^[28]^ **(Supplementary Table 1)**. To quantify the aNK phenotype across NK subclusters, we calculated an aNK module score and compared its distribution among clusters. The aNK population displayed significantly higher module scores than other NK subsets, supporting its identity as a transcriptionally distinct NK state within the dataset **(Fig. 1C)**. We then examined how NK subset composition changed across CRT time points. Notably, the relative frequency of aNK cells increased following CRT, with enrichment already apparent after the first fraction and persisting after the second fraction **(Fig. 1D)**, indicating a rapid and sustained therapy-associated shift in NK cell composition.

**Figure 1.**
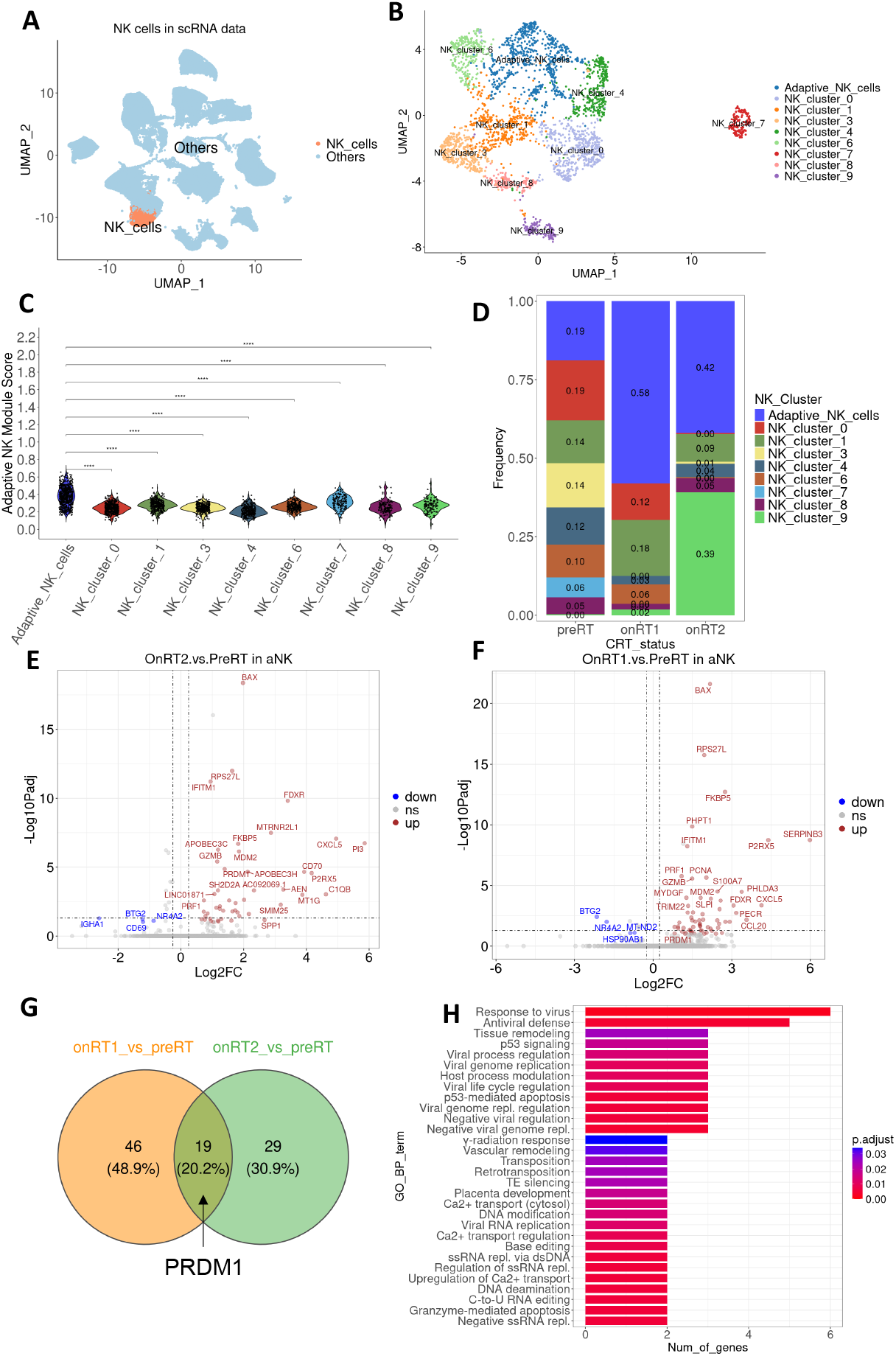
CRT induces upregulation of the transcription factor PRDM1 in aNK cells. **(A)** UMAP visualization shows NK cells separated from the total cells. **(B)** UMAP visualization of NK cell subclusters derived from the extracted NK cell population.**(C)** Violin plot displaying the distribution of aNK cell module scores across all clusters, with pairwise Mann-Whitney-Wilcoxon tests performed using aNK cluster as the reference group. Statistical significance is denoted by: **** *p* < 0.0001. **(D)** Stacked bar plots show the relative frequencies of NK subsets in pre-treatment (preRT), first-time CRT (onRT1), and second-time CRT (onRT2). **(E-F)** Volcano plots depict differentially expressed genes in aNK cells comparing onRT2 vs preRT **(E)** and onRT1 vs preRT **(F). (G)** A Venn diagram highlights shared upregulated genes between the two comparisons, including PRDM1. **(H)** Gene Ontology (GO) analysis of overlapping genes reveals enriched biological processes.

To characterize transcriptional remodeling of aNK cells during CRT, we performed differential expression analyses comparing **onRT2 vs preRT** and **onRT1 vs preRT** within the aNK compartment. Both comparisons revealed broad CRT-associated transcriptional changes, including induction of cytotoxic and interferon/virus-response-related genes **(Fig. 1E–F)**. A set of upregulated genes was shared between onRT1 and onRT2 relative to preRT, suggesting a conserved CRT-induced aNK program across treatment stages **(Fig. 1G)**. Importantly, **PRDM1** emerged as a consistently upregulated transcription factor in both comparisons with high expression in aNK cell cluster **(Supplementary Fig.1A)**. Gene Ontology (GO) enrichment analysis of overlapping upregulated genes highlighted immune activation and antiviral-response pathways consistent with enhanced effector programs in CRT-associated aNK cells **(Fig. 1H)**. Together, these results indicate that CRT is accompanied by enrichment of aNK cells and coordinated activation of cytotoxic/antiviral transcriptional programs, with PRDM1 as a recurrently induced regulator in aNK cells.

### Pseudotime analysis reveals dynamic state transitions and PRDM1 induction along the CRT-associated NK trajectory

To further characterize the CRT-associated transcriptional dynamics within NK cells, we performed pseudotime trajectory analysis on the NK cell compartment. Mapping cells onto the UMAP embedding with inferred pseudotime values revealed an ordered continuum spanning multiple NK states, suggesting progressive transcriptional transitions, in which aNK cell cluster located at one terminal state **(Fig. 2A)**. Along the trajectory, cells from onRT1 and onRT2 were preferentially distributed toward terminal pseudotime states **(Fig. 2B)**.

**Figure 2.**
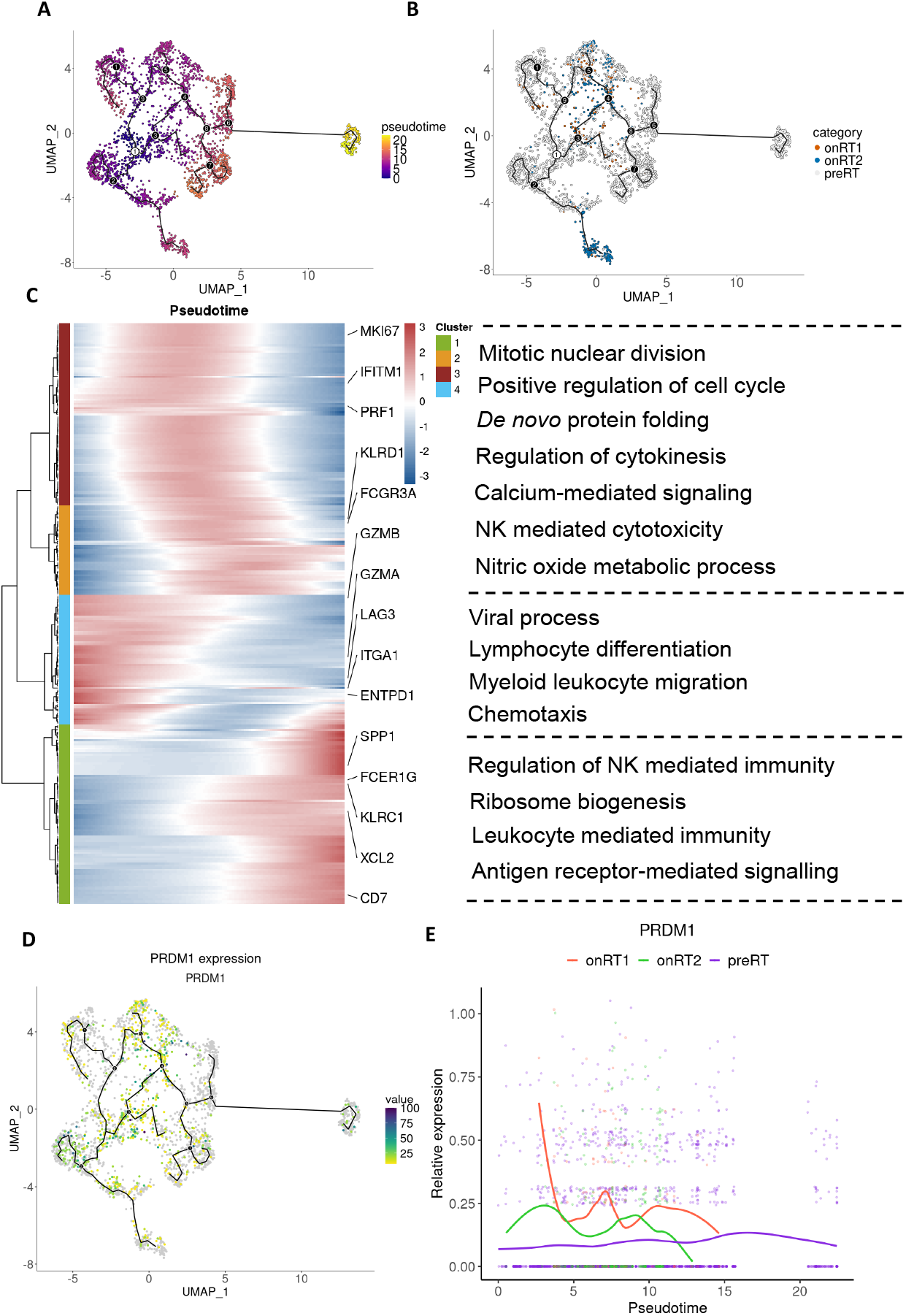
Pseudotime trajectory analysis reveals dynamic transcriptional changes of NK cells. **(A)** UMAP embedding overlaid with pseudotime values illustrates the inferred developmental trajectory of NK cells, with color indicating progression along pseudotime. **(B)** Cells are colored by treatment condition (preRT, onRT1, and onRT2) and projected onto the trajectory to show their distribution along the developmental path. **(C)** Heatmap displays the modules of dynamically expressed genes along pseudotime, with representative genes labeled and major enriched GO biological processes summarized on the right. **(D)** UMAP visualization of PRDM1 expression demonstrates its spatial distribution along the trajectory, with color intensity indicating expression levels. **(E)** Scatter plot shows PRDM1 expression dynamics across pseudotime in preRT, onRT1, and onRT2 conditions, with fitted curves indicating overall trends.

Consistent with this progression, genes that varied dynamically along pseudotime organized into modules with coordinated expression patterns, capturing distinct phases of NK cell reprogramming **(Fig. 2C)**. Early-to-intermediate pseudotime segments were characterized by enrichment of processes related to cell cycle and protein folding, whereas later segments were associated with immune functional programs, including cytokine regulation, NK-mediated cytotoxicity, and antiviral response pathways **(Fig. 2C)**. This pattern suggests a progressive functional maturation of NK cells along pseudotime, marked by the distinct modules with varied patterns of gene expressions.

Given that PRDM1 was consistently induced in aNK cells after CRT in differential expression analyses, we next examined PRDM1 expression along the inferred NK trajectory. PRDM1 expression was spatially enriched along specific regions of the trajectory on UMAP **(Fig. 2D)** and displayed stage-dependent expressions across pseudotime **(Fig. 2E)**. PRDM1 expression exhibited a pseudotime-dependent pattern and differed by treatment stage. Compared with preRT, on-treatment cells (onRT1 and onRT2) showed higher PRDM1 expression over early-to-intermediate pseudotime, with a pronounced increase in onRT1. In contrast, preRT cells maintained relatively low and stable PRDM1 levels across the trajectory, indicating that PRDM1 upregulation is associated with the treatment-enriched NK states along pseudotime **(Fig. 2E)**. Together, these analyses indicate that CRT is accompanied by a directional shift in NK cell states along pseudotime and that PRDM1 induction occurs within this trajectory.

### GRN analysis identifies PRDM1 as a hub transcriptional regulator associated with aNK cells and CRT

To identify key transcriptional regulators of aNK cell programs and to assess how these regulatory networks change during CRT, we performed gene regulatory network (GRN) analysis and ranked transcription factors by **betweenness centrality**, a measure of network “hubness” reflecting how strongly a node connects multiple regulatory modules. In the GRN constructed from aNK cells, centrality ranking highlighted a core set of regulatory hubs dominated by immediate-early and immune-associated transcription factors, including FOS, KLF2, KLF6, IRF1, and others. Notably, PRDM1 also appeared among the top-ranked transcription factors, indicating that it occupies a structurally important position within the aNK regulatory network **(Fig. 3A)**.

**Figure 3.**
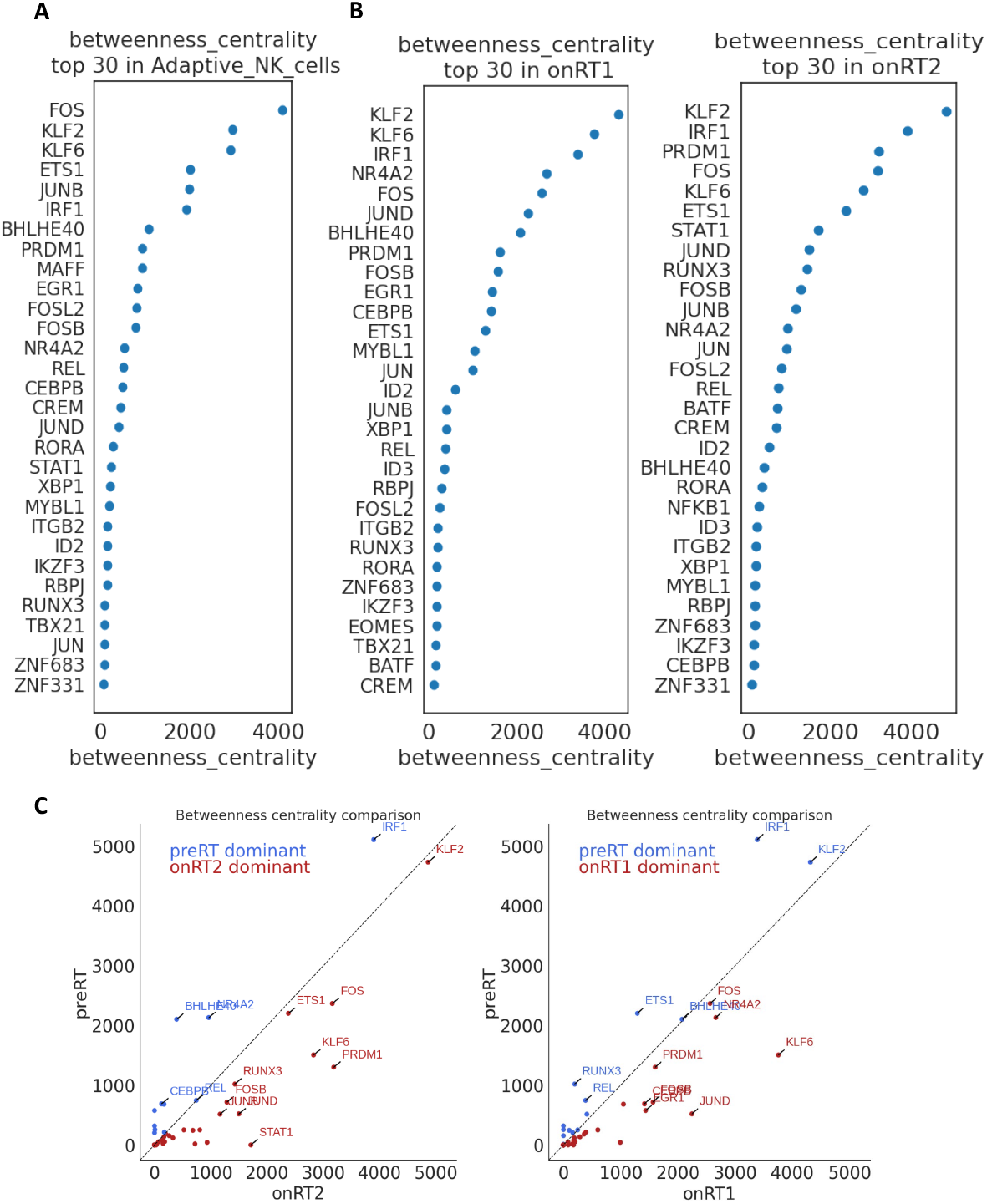
Gene regulatory network (GRN) analysis reveals key transcriptional regulators in aNK cells during CRT. **(A-B)** Dot plots show the top 30 transcription factors ranked by betweenness centrality in GRNs constructed from aNK cells **(A)**, onRT1 **(B, left)**, and onRT2 **(B, right). (C)** Scatter plots compare betweenness centrality values between preRT and onRT2 **(C, left)** and between preRT and onRT1 **(C, right)**, where genes above the diagonal represent preRT-dominant regulators and those below represent onRT-dominant regulators.

We next assessed whether the regulatory architecture was altered across CRT stages by constructing condition-specific GRNs for on-treatment samples. Betweenness centrality ranking of the onRT1 and onRT2 GRNs showed a reordered set of hub regulators compared with baseline **(Fig. 3B)**. Notably, PRDM1 consistently remained among the top-ranked transcription factors in both onRT1 and onRT2, indicating that it occupies a central position in the treatment-associated network architecture in addition to being transcriptionally induced **(Fig. 3B)**. PRDM1 was likewise prioritized by other centrality metrics, including degree and eigenvector centrality **(Supplementary Fig 2A-B)**.

To further quantify treatment-related changes in network structure, we directly compared transcription factor betweenness centrality values between preRT and on-treatment states. These comparisons identified PRDM1 as one of the regulators with increased centrality after treatment (onRT-dominant) and those with higher centrality at baseline (preRT-dominant) **(Fig. 3C)**. Together, these results indicate that CRT is accompanied by measurable rewiring of the aNK GRN and highlight PRDM1 as a recurrent, treatment-associated hub regulator across onRT1 and onRT2 states.

### CellOracle *in silico* perturbation predicts PRDM1-dependent trajectory dynamics in NK cells and aNK cells

Given the induction and network centrality of PRDM1 in CRT-associated aNK cells, we next used CellOracle^[34]^ to perform *in silico* perturbation and evaluate how PRDM1 disruption reshapes inferred cell-state dynamics. Under control conditions, developmental flow vectors overlaid on the NK UMAP revealed coherent directional transitions across NK cell states, consistent with an ordered progression in the NK compartment **(Fig. 4A)**. Simulating PRDM1 knockout (KO) produced a marked change in the predicted direction and magnitude of cell identity shifts across the embedding **(Fig. 4B)**, indicating that PRDM1 contributes to the inferred dynamics of NK state transitions.

**Figure 4.**
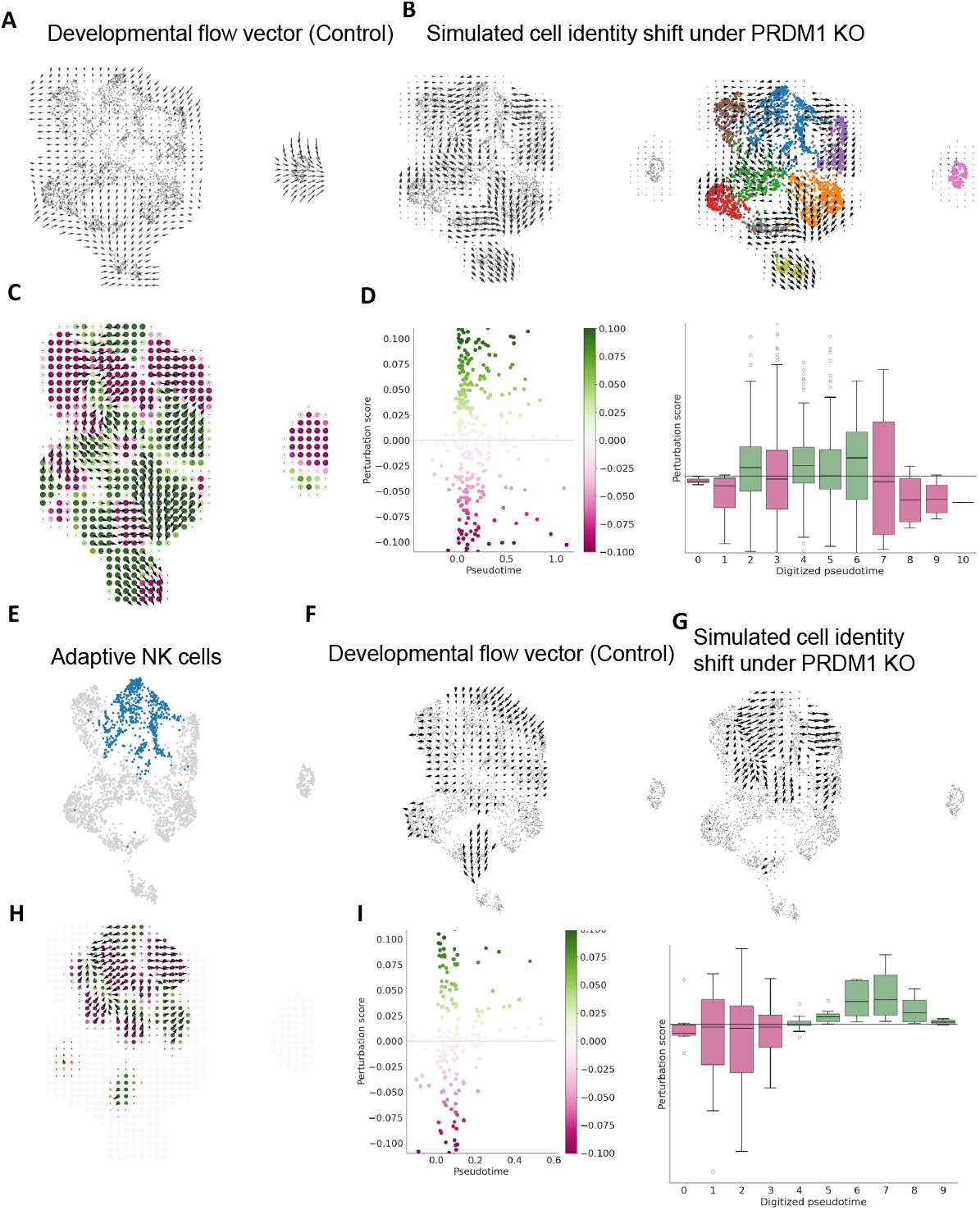
CellOracle-based *in silico* knockout (KO) of PRDM1 predicts altered developmental trajectories of aNK cells. **(A)** Developmental flow vectors inferred under control conditions illustrate the baseline directional dynamics of NK cell states. **(B)** Simulated cell identity shifts following PRDM1 in silico knockout (KO) reveal marked alterations in predicted cell state transitions across the UMAP space. **(C)** Perturbation score (PS) distribution projected onto the NK cell UMAP, where positive (green) values indicate promoted differentiation and negative (purple) values indicate blocked differentiation based on the inner product between perturbation and developmental flow vectors. **(D)** PS along pseudotime are visualized using two complementary approaches: a scatter plot showing continuous PS distribution across pseudotime (left) and box plots summarizing PS across discretized pseudotime bins (right). **(E)** UMAP visualization highlighting aNK cells within the NK cell population. **(F-G)** Developmental flow vectors under control conditions **(F)** and simulated cell identity shifts following PRDM1 in silico knockout **(G)**, displayed specifically for the aNK cell cluster. **(H)** (PS) distribution projected onto the aNK cell cluster on UMAP. **(I)** PS dynamics along pseudotime within the aNK cluster, shown as a scatter plot across continuous pseudotime (left) and summarized by box plots across discretized pseudotime bins (right).

To quantify perturbation effects relative to the baseline developmental flow, we computed a perturbation score (PS) and projected it onto the NK UMAP. PS was defined as the inner product (dot product) between the 2D perturbation-simulation vector and the baseline developmental-flow vector at each location, capturing both vector direction and magnitude; thus, positive PS indicates alignment with the developmental flow (predicted promotion of differentiation), whereas negative PS indicates opposition to the flow (predicted blockade of differentiation) **(Fig. 4C)**. When PS was examined along pseudotime, PRDM1 KO showed pseudotime-dependent effects on total NK dynamics, with the strongest contradictory impact at late pseudotime where PRDM1 perturbation was associated with the opposite direction to the natural developmental flow **(Fig. 4D)**.

We then focused on the aNK compartment highlighted on the UMAP **(Fig. 4E)**. Within the aNK cluster, baseline developmental flow vectors remained directional **(Fig. 4F)**, whereas PRDM1 KO simulation altered the predicted identity shift patterns **(Fig. 4G)**. Consistently, PS projected onto the aNK UMAP demonstrated localized regions of positive and negative perturbation effects **(Fig. 4H)**, and Pseudotime analysis restricted to aNK cells showed a clear shift in PRDM1 KO effects across the continuum, with negative PS values at early pseudotime indicating opposition to the developmental flow (predicted blockade of differentiation) and positive PS values at later pseudotime indicating alignment with the flow (predicted promotion of differentiation) **(Fig. 4I)**. Together, these CellOracle simulations support a model in which PRDM1 contributes to maintaining the inferred developmental flow and state-transition dynamics of NK cells, with pronounced and stage-dependent effects within the aNK compartment.

### scTenifoldKnk virtual KO of PRDM1 predicts broad downstream network perturbation with prominent metabolic signatures in aNK cells

To further define downstream programs regulated by PRDM1 in aNK cells, we performed ***in silico*** PRDM1 KO using **scTenifoldKnk**^[35]^ and quantified gene-level perturbation effects. The resulting perturbed gene set was mapped onto a PRDM1-centered regulatory context, visualized in Cytoscape^[36]^, highlighting PRDM1-associated connectivity among affected targets and their relative perturbation strength **(Fig. 5A)**. Gene-level perturbation statistics showed a subset of genes exhibiting strong deviation upon PRDM1 virtual KO, as summarized by Z-scores and significance in the volcano plot **(Fig. 5B)**, indicating widespread transcriptional consequences predicted from PRDM1 disruption.

**Figure 5.**
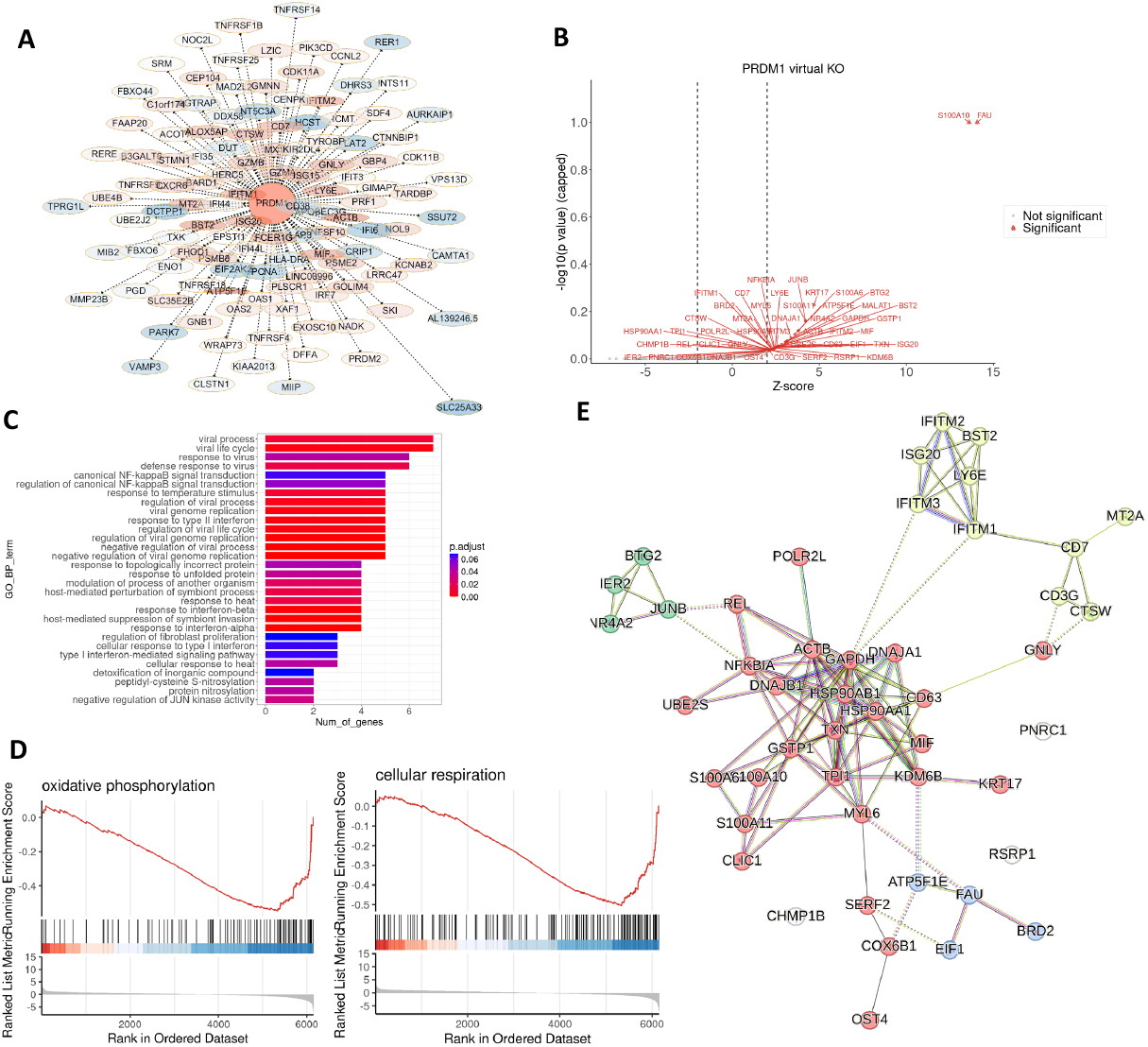
scTenifoldKnk-based *in silico* KO of PRDM1 reveals downstream regulatory and functional effects in aNK cells. **(A)** Gene regulatory network (GRN) visualization generated in Cytoscape highlights PRDM1-centered perturbed genes, with nodes representing affected genes and colored according to their Z-scores. (B) Volcano plot shows genes most affected by PRDM1 virtual KO, with significantly perturbed genes highlighted. **(C)** Bar plot showing GO enrichment results of the top genes affected by PRDM1 virtual KO. **(D)** Gene set enrichment analysis (GSEA) demonstrates significant enrichment of oxidative phosphorylation and cellular respiration pathways upon PRDM1 perturbation. **(E)** Protein–protein interaction (PPI) network of the top genes affected by PRDM1 virtual knockout, constructed using STRING and visualized to show functional connectivity among perturbed genes.

We next asked which biological processes were overrepresented among the top perturbed genes. GO enrichment analysis revealed that the scTenifoldKnk-derived gene set was strongly enriched for immune response-related programs, particularly antiviral/defense-associated processes (**Fig. 5C)**. In parallel, GSEA of the scTenifoldKnk network Z score ranked gene list highlighted a dominant mitochondrial bioenergetic signature, with enriched GO terms related to electron transport chain activity, respiratory complex assembly, and ATP synthesis coupled transport, and concordant KEGG enrichment for oxidative phosphorylation **(Supplementary Fig.3A-B)**. Importantly, oxidative phosphorylation and cellular respiration showed negative enrichment, indicating that PRDM1 virtual knockout shifts these two pathways toward the negative end of the ranked perturbation profile, inferring the disrupted mitochondrial metabolism related to PRDM1 deletion **(Fig. 5D)**. Finally, following scTenifoldKnk-based PRDM1 virtual knockout, we constructed a protein–protein interaction (PPI) network using STRING and performed network clustering, which resolved four densely connected modules among the top perturbed genes **(Fig. 5E)**. The first module comprised canonical interferon-stimulated/antiviral factors, including IFITM1/2/3, ISG20, BST2, and LY6E, forming a tightly interconnected cluster consistent with coordinated regulation of antiviral programs. A second module captured cytotoxic and lymphocyte effector features, with CD7, CD3G, CTSW, and GNLY connected to additional stress-associated components (e.g., MT2A), indicating perturbation of effector-associated circuitry. A third module was enriched for immediate-early and inflammatory transcriptional regulators, including REL, NFKBIA, JUNB, NR4A2, BTG2, IER2, and UBE2S, consistent with rewiring of NF-κB/AP-1–linked response networks upon PRDM1 perturbation. The fourth module was dominated by metabolic and proteostasis-related genes, centered on chaperones and redox/stress proteins such as HSP90AA1/HSP90AB1, DNAJA1/DNAJB1, TXN, GSTP1, GAPDH, ACTB, TPI1, and MIF, and extending to mitochondrial/translation-associated components including ATP5F1E, COX6B1, EIF1, and FAU, supporting coordinated disruption of bioenergetic and protein homeostasis pathways. Notably, several connector nodes bridged these modules, indicating that PRDM1 perturbation is predicted to jointly impact antiviral signaling, inflammatory regulation, cytotoxic effector programs, and metabolic/proteostasis networks within aNK cells.

## Methods

### Data source, preprocessing, and NK cell subclustering

Single-cell RNA-seq data from cervical cancer patients were obtained from GEO (GSE297041) and analyzed using a preprocessed Seurat object saved as an RDS file. NK cells were extracted from the global dataset based on the cell identity label assigned by the cell annotation. For preprocessing within the NK compartment, mitochondrial genes were defined by the MT-prefix and the mitochondrial transcript fraction was calculated per cell; expression values were then normalized, 3,000 highly variable genes were selected, and the data were scaled with regression of mitochondrial content. Dimensionality reduction was performed by PCA, and the NK subset was subclustered by constructing a nearest-neighbor graph using the first 17 principal components followed by graph-based clustering at resolution 0.5. The resulting NK subclusters were visualized using a UMAP embedding generated from the same principal components. All the R processing were performed using Seurat (v5.3.0)^[37]^.

### Cell-type annotation of NK cells and identification of aNK cells

Cell-type identities in the full cervical cancer scRNA-seq dataset were assigned using SingleR (v2.10.0)^[38]^. Briefly, the raw count matrix was queried against curated human cell atlas references, and each cell was labeled by the best-matching immune or primary cell-type profile; these predicted labels were then visualized on the global UMAP and summarized across unsupervised clusters. To derive a robust cluster-level annotation, we computed the contingency table between predicted labels and clustering assignments and assigned each cluster the most frequent predicted label, which was then propagated back to all cells as a cluster annotation.

For finer annotation within the NK compartment and specific identification of aNK cells, we applied a marker-based scoring approach using scType^[31]^. Positive and negative marker gene sets were scored on the scaled expression matrix of the NK subset, and scores were summarized at the cluster level to determine the best-supported NK subtype for each cluster. The subtype with the highest aggregated score was assigned as the annotation, while clusters with low-confidence support, defined as a top score below 25% of the cluster cell number, were labeled as Unknown.

### Differential expression and enrichment analyses

aNK cells were extracted from the NK subset based on the assigned NK subtype annotation and grouped by treatment stage (preRT, onRT1, onRT2). Differential expression was performed within the aNK compartment by comparing onRT1 versus preRT and onRT2 versus preRT using a Wilcoxon rank-sum test, including genes detected in at least 10% of cells in either group. For visualization and downstream summaries, genes were categorized as upregulated or downregulated using an adjusted P value threshold of 0.1 together with an absolute log2 fold-change cutoff of 0.5, and volcano plots were generated with selected significant genes labeled (including PRDM1).

To identify conserved treatment-associated induction signatures, upregulated gene sets from onRT1 versus preRT and onRT2 versus preRT were intersected and visualized using set-overlap plots. Functional enrichment analysis was then performed on the overlapping upregulated genes using GO biological process (GO-BP) over-representation testing with Benjamini–Hochberg correction, applying significance thresholds of adjusted P < 0.01 and q < 0.05; enriched terms were ranked by adjusted P value and visualized using bar plots of the top pathways.

### Pseudotime trajectory analysis

Trajectory inference was performed on the NK cell subset using monocle3 (v1.4.26)^[39]^ single-cell pseudotime framework. The count matrix and corresponding cell metadata from the NK subset were converted into a trajectory object, followed by dimensionality reduction and construction of a UMAP embedding. To ensure consistency with the clustering-based visualization used throughout the study, the trajectory UMAP coordinates were replaced with the UMAP embedding previously computed in the Seurat NK analysis. A principal graph was then learned on the UMAP space (without partitioning) and cells were ordered along the graph to assign pseudotime values, using both default ordering and an automated root-node selection strategy based on the earliest principal-graph node. Pseudotime and treatment stage were visualized on the trajectory, and PRDM1 expression was projected onto the same embedding. To identify genes with significant spatial dependence along the trajectory, graph-based tests were performed and significant genes (q value < 0.05) were grouped into co-expression modules; module-level expression patterns were summarized across NK cell types and visualized as heatmaps. For higher-level interpretation, pseudotime-ordered expression matrices of significant genes were smoothed, z-scored, binned along pseudotime, and clustered into four major gene modules, followed by GO biological process enrichment analysis for each module using multiple-testing correction.

### Gene regulatory network inference and centrality analysis

GRNs were inferred from the NK-cell AnnData object using CellOracle with a human promoter-based prior GRN as the base reference. Raw (unscaled) count matrices were used as input, and cells were grouped either by NK subtypes to generate subtype-specific GRNs or by treatment stage (category; preRT, onRT1, onRT2) to generate condition-specific GRNs. After dimensionality reduction and kNN-based imputation, GRN edges were estimated within each group and filtered using a stringent significance cutoff (p < 0.001), retaining the top-ranked edges by absolute coefficient (2,000 links). For each inferred GRN, network topology metrics were computed and transcription factors were ranked by centrality-based network scores, including betweenness centrality, degree centrality, and eigenvector centrality, to prioritize hub regulators. Centrality rankings were visualized as score-rank plots within aNK and within each CRT stage, and treatment-associated network rewiring was assessed by pairwise comparison of transcription factor betweenness centrality between on-treatment (onRT1 or onRT2) and baseline (preRT) networks, highlighting regulators with stage-enriched connectivity.

### CellOracle *in silico* transcription factor perturbation

To assess the predicted functional impact of PRDM1 on NK-state dynamics, we performed *in silico* transcription factor perturbation using CellOracle based on the inferred NK GRNs. Briefly, the fitted GRN model was used to simulate PRDM1 knockout by setting PRDM1 activity to zero and propagating the perturbation through the network (three propagation steps). Transition probabilities were then estimated using a large kNN neighborhood (200 neighbors) and projected onto the UMAP embedding to calculate cell-identity shift vectors, which were visualized as vector fields on a grid. In parallel, a developmental reference flow was computed from pseudotime using a gradient-based approach and similarly represented as a grid-based vector field. To quantify the relationship between the perturbation-induced shift and the baseline developmental flow, we calculated a perturbation score (PS) as the inner product between the two 2D vectors at each grid location, capturing both directionality and magnitude; positive values indicate alignment with the developmental flow, whereas negative values indicate opposition. Perturbation scores were visualized on the NK UMAP and further summarized across pseudotime using both continuous representations and digitized pseudotime bins. The same workflow was repeated after restricting the analysis to cells within the aNK cluster to resolve PRDM1-dependent effects specifically along the aNK pseudotime continuum.

### scTenifoldKnk virtual knockout and downstream analyses

To model PRDM1-dependent regulatory effects within aNK cells, we performed virtual KO using scTenifoldKnk on the aNK count matrix. aNK cells were subsetted from the NK object, and the count matrix was extracted for downstream network inference. For quality control and to reduce technical dominance from highly abundant housekeeping features, genes matching ribosomal, mitochondrial, and histone-related patterns were optionally removed prior to saving a filtered expression matrix for sensitivity analyses. Virtual knockout was then performed by specifying PRDM1 as the perturbed gene and running the network construction using multiple subsampled cell sets and repeated network reconstructions (500 cells per subsample, 10 networks, 30 components, multi-core execution). The resulting differential regulation output was ranked by significance, and genes with strong perturbation effects were defined using a Z-score threshold (>2) while excluding PRDM1 itself. Perturbation effects were visualized using a volcano-style plot based on Z-scores and −log10(*P*) values with significant genes labeled.

Downstream functional interpretation was carried out using both over-representation and rank-based enrichment approaches. Over-representation analysis was performed on the significantly affected genes using GO biological process enrichment with multiple-testing correction (Benjamini–Hochberg) and stringent cutoffs (adjusted *P* < 0.01; *q* < 0.05). In parallel, gene set enrichment analysis was performed using the full gene list ranked by network Z-scores, including GO-BP GSEA and KEGG GSEA with standard gene-set size constraints and a significance cutoff of *P* < 0.05, and representative enriched pathways were visualized using enrichment plots.

### Cytoscape subnetwork construction

To visualize a PRDM1-centered differential regulatory subnetwork, we extracted the inferred regulatory adjacency matrices from the wild-type and PRDM1-KO network reconstructions and computed the KO-minus-WT difference matrix. PRDM1 outgoing and incoming differential edges were ranked by absolute weight, and a neighborhood of top PRDM1-connected genes was selected to define a focused subnetwork. Differential edges within this subnetwork were then converted into an edge table containing source, target, signed weight and absolute weight, and a node table was generated by integrating scTenifoldKnk gene-level statistics (Z-scores, *P* values, and significance calls) for the same set of nodes. The cleaned edge and node tables were exported as CSV files for downstream visualization and styling in Cytoscape software.

### Code availability

All code used for data processing, analysis, and figure generation in this study is publicly available at: https://github.com/yizhesuncode/Cervical_RT_aNK_codes/.

## Discussion

In this study, we reanalyzed a longitudinal cervical cancer scRNA-seq dataset collected before CRT and during treatment and observed a reproducible expansion of aNK cells together with induction of antiviral and cytotoxic transcriptional programs. These changes are consistent with the established ability of radiotherapy and chemotherapy to remodel the tumor immune microenvironment by promoting immunogenic cell death and DAMP release, which can enhance dendritic-cell activation and antigen presentation, and by engaging inflammatory cytokine and interferon pathways that reshape immune-state composition. In parallel, irradiation can increase tumor-cell immunogenicity through increased MHC class I antigen presentation and stress-associated changes that influence susceptibility to cytotoxic lymphocytes, including NK cells^[6]^. Together, these therapy-conditioned signals provide a plausible mechanistic context for the shift toward more differentiated or functionally primed NK states during on-treatment sampling in cervical cancer.

A key biological contribution of our work is the identification of PRDM1 as a CRT-associated transcriptional regulator tightly linked to the aNK compartment. PRDM1 (Blimp-1) is a well-established controller of lymphocyte differentiation programs, and prior work in human NK cells has shown that PRDM1 can regulate effector cytokine outputs and broader activation restraint, supporting a direct role in shaping NK functional states^[40]^. Consistent with this, our GRN centrality analyses suggest that PRDM1 is not only transcriptionally induced during on-treatment states but also embedded as a highly connected node within treatment-associated regulatory architecture, arguing that PRDM1 may integrate therapy-driven inflammatory cues into coordinated downstream programs in NK cells.

Our findings also align PRDM1 with the broader biology of immune memory and tumor-infiltrating adaptive-like NK states. Beyond the current cervical CRT setting, PRDM1 has been highlighted as a key transcriptional regulator in tumor-infiltrating aNK cells in ovarian cancer, supporting its potential cross-tumor relevance for aNK regulation^[41]^. In addition, recent tumor-immune surveillance work supports a functional requirement for Prdm1 in group 1 ILCs in cancer contexts^[42]^, consistent with the idea that PRDM1 contributes to maintaining functional NK-state heterogeneity under tumor-associated stress.

Finally, our *in silico* perturbation analyses provide a functional framework for interpreting PRDM1 association with CRT-enriched aNK states. CellOracle suggested that PRDM1 influences inferred NK state-transition dynamics in a pseudotime-dependent manner, while scTenifoldKnk indicated broad downstream network consequences spanning immune-response modules and metabolism. Notably, oxidative phosphorylation and cellular respiration showed negative enrichment in GSEA, consistent with coordinated suppression of mitochondrial respiratory programs upon PRDM1 perturbation. Together with STRING-based PPI clustering, these results support a model in which PRDM1 coordinates coupled antiviral and inflammatory circuitry with cytotoxic and bioenergetic or proteostasis programs in aNK cells.

There are important limitations. First, mechanistic inferences here are derived from computational perturbation and network reconstruction and therefore require experimental validation. Second, scRNA-seq captures transcriptional snapshots and does not directly measure protein activity, chromatin state, or post-transcriptional regulation, which are central to PRDM1 function. Third, treatment timing and patient heterogeneity, including HPV biology, potential CMV status, prior immune history, and regimen variability, can influence NK states and are typically incompletely captured in public datasets. Finally, aNK biology is classically linked to HCMV-driven differentiation with epigenetic remodeling, and the extent to which tumor-infiltrating aNK cells in cervical cancer represent canonical HCMV-associated aNK versus convergent adaptive-like differentiation remains an open question.

Despite these limitations, our study links CRT-associated NK-state remodeling to a PRDM1-centered regulatory axis in aNK cells. Integrating longitudinal single-cell profiling with regulatory-network inference and *in silico* perturbation suggests that PRDM1 is associated with aNK enrichment and may coordinate antiviral and cytotoxic transcriptional programs with mitochondrial respiratory pathways in the CRT-conditioned tumor microenvironment. Collectively, these results refine the transcriptional framework of therapy-driven NK adaptation in cervical cancer and highlight PRDM1 as a central node within this response.

## Author contributions

M.W. and T.Z. contributed equally to this work. Y.S. conceived and supervised the study. M.W., T.Z., and Y.S. designed the research strategy and analytical framework. M.W. and T.Z. collected and curated the clinical and/or sequencing data and performed primary data processing. M.W. performed advanced computational analyses, integrative bioinformatics, visualization, and interpretation of results. W.S. and J.S. contributed to data organization, annotation, and supporting analyses. W.Z. provided clinical expertise and assisted with result interpretation. M.W., T.Z., and Y.S. drafted the manuscript. All authors reviewed, edited, and approved the final manuscript, and agree to be accountable for all aspects of the work.

---

**Supplementary Figure 1.**
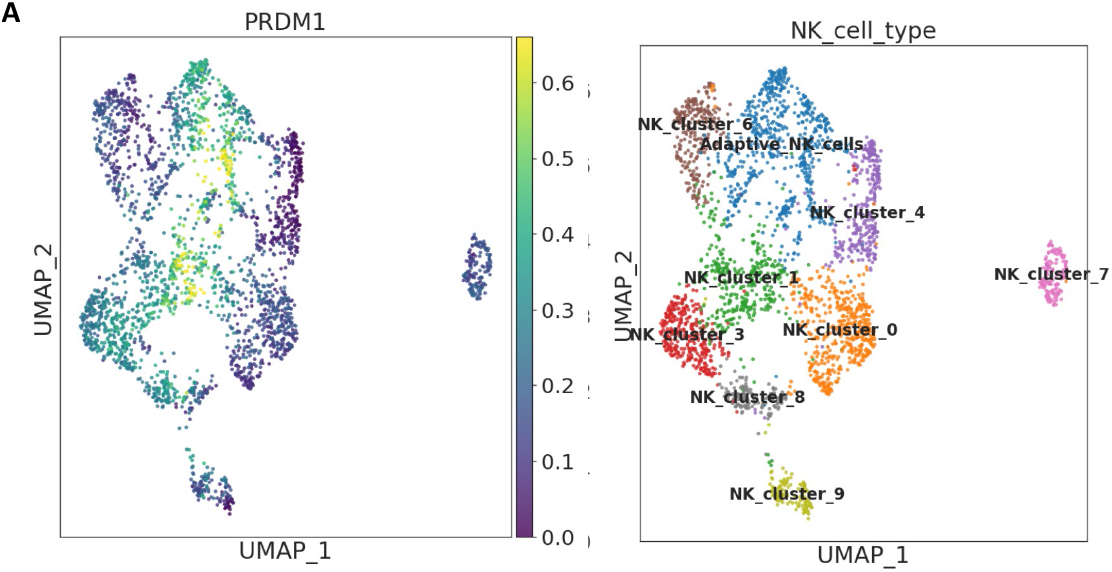
PRDM1 is preferentially expressed in aNK cells. **(A)** UMAP projection of NK cells colored by PRDM1 expression (left) and annotated by NK cell subtypes (right).

**Supplementary Figure 2.**
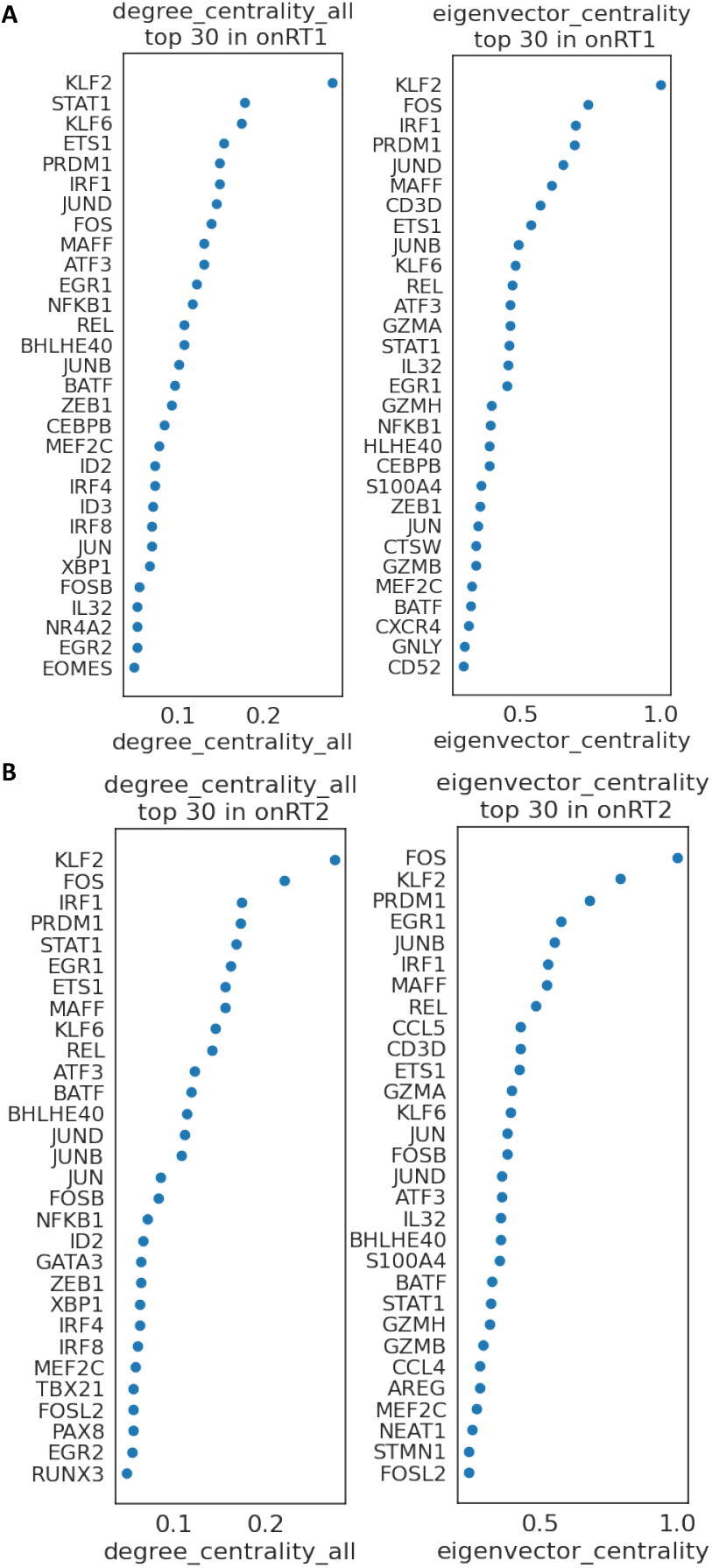
PRDM1 ranks among the top hub regulators in both onRT1 and onRT2 networks. **(A-B)** Dot plots show the top 30 transcription factors ranked by degree and eigenvector centrality in GRNs constructed from onRT1 **(A)**, and onRT2 **(B)**.

**Supplementary Figure 3.**
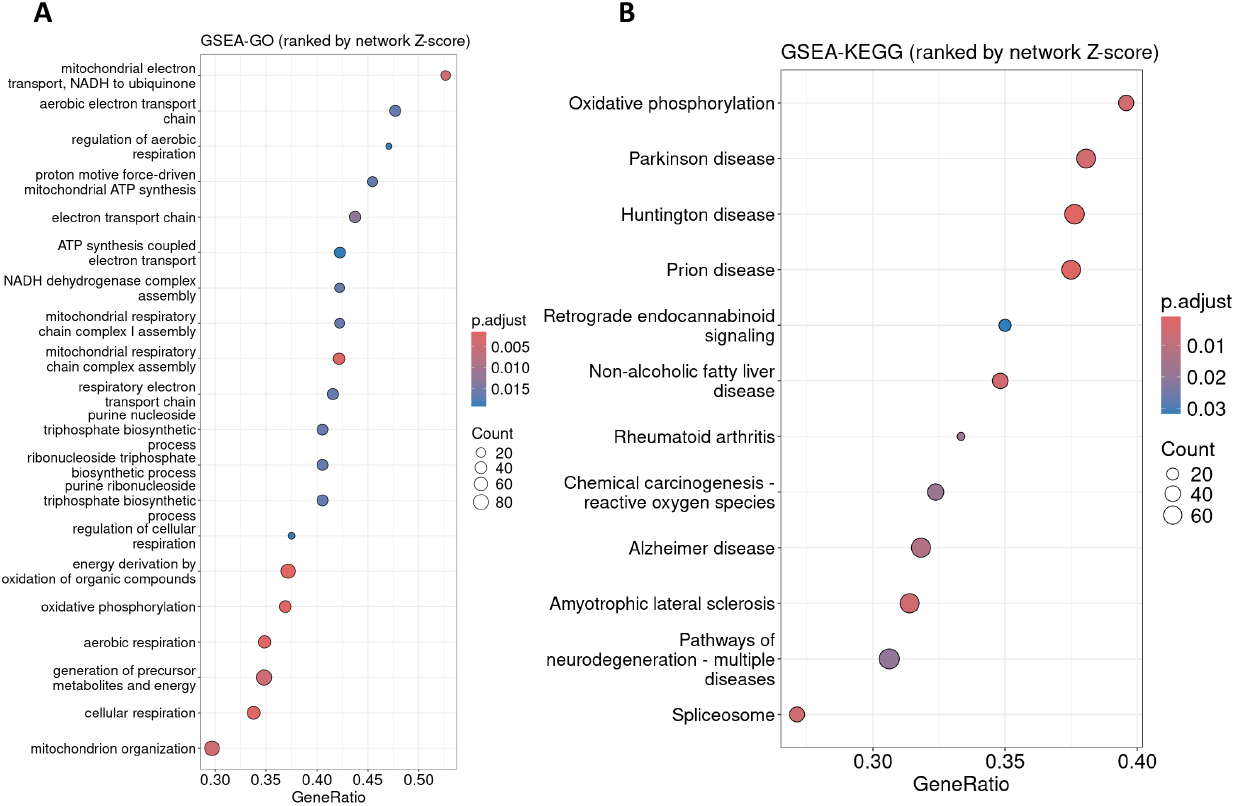
Metabolic and mitochondrial pathways are enriched among genes perturbed by PRDM1 virtual KO. **(A-B)** Bubble plot showing GO **(A)** enrichment and KEGG**(B)** results for the gene set identified by scTenifoldKnk following PRDM1 virtual knockout. Gene sets are ranked by the network Z-score derived from scTenifoldKnk.

