## Supplementary Table 1 for "PRDM1 Drives Chemoradiotherapy-associated Enrichment of Adaptive NK Cells in Cervical Cancer"

**Supplementary Table 1. Gene signatures used for aNK subcluster annotation**

| CellType | tissueType | cellName | geneSymbolmore1(Positively expressed) | geneSymbolmore2 (Negatively expressed) |
|---|---|---|---|---|
| 0 | Immune system | Adaptive_NK | FCGR3A | KLRC1,FCER1G,SYK |
| 0 | Immune system | Adaptive_NK | FGFBP2 | KLRC1,FCER1G,SYK |
| 0 | Immune system | Adaptive_NK | KLRF1 | KLRC1,FCER1G,SYK |
| 0 | Immune system | Adaptive_NK | KLF2 | KLRC1,FCER1G,SYK |
| 0 | Immune system | Adaptive_NK | PLAC8 | KLRC1,FCER1G,SYK |
| 0 | Immune system | Adaptive_NK | GZMB | KLRC1,FCER1G,SYK |
| 0 | Immune system | Adaptive_NK | GZMH | KLRC1,FCER1G,SYK |
| 0 | Immune system | Adaptive_NK | CX3CR1 | KLRC1,FCER1G,SYK |
| 0 | Immune system | Adaptive_NK | ARL4C | KLRC1,FCER1G,SYK |
| 0 | Immune system | Adaptive_NK | PRF1 | KLRC1,FCER1G,SYK |
| 0 | Immune system | Adaptive_NK | EFHD2 | KLRC1,FCER1G,SYK |
| 0 | Immune system | Adaptive_NK | CYBA | KLRC1,FCER1G,SYK |
| 0 | Immune system | Adaptive_NK | NKG7 | KLRC1,FCER1G,SYK |
| 0 | Immune system | Adaptive_NK | PLEK | KLRC1,FCER1G,SYK |
| 0 | Immune system | Adaptive_NK | S1PR5 | KLRC1,FCER1G,SYK |
| 0 | Immune system | Adaptive_NK | CST7 | KLRC1,FCER1G,SYK |
| 0 | Immune system | Adaptive_NK | ZEB2 | KLRC1,FCER1G,SYK |
| 0 | Immune system | Adaptive_NK | DTHD1 | KLRC1,FCER1G,SYK |
| 0 | Immune system | Adaptive_NK | SPON2 | KLRC1,FCER1G,SYK |
| 0 | Immune system | Adaptive_NK | ADGRG1 | KLRC1,FCER1G,SYK |
| 0 | Immune system | Adaptive_NK | RIPOR2 | KLRC1,FCER1G,SYK |
| 0 | Immune system | Adaptive_NK | S100A4 | KLRC1,FCER1G,SYK |
| 0 | Immune system | Adaptive_NK | TGFBR3 | KLRC1,FCER1G,SYK |
| 0 | Immune system | Adaptive_NK | PRSS23 | KLRC1,FCER1G,SYK |
| 0 | Immune system | Adaptive_NK | IGFBP7 | KLRC1,FCER1G,SYK |
| 0 | Immune system | Adaptive_NK | GZMM | KLRC1,FCER1G,SYK |
| 0 | Immune system | Adaptive_NK | TTC38 | KLRC1,FCER1G,SYK |
| 0 | Immune system | Adaptive_NK | PFN1 | KLRC1,FCER1G,SYK |
| 0 | Immune system | Adaptive_NK | CD247 | KLRC1,FCER1G,SYK |
| 0 | Immune system | Adaptive_NK | CEP78 | KLRC1,FCER1G,SYK |
| 0 | Immune system | Adaptive_NK | LITAF | KLRC1,FCER1G,SYK |
| 0 | Immune system | Adaptive_NK | ACTB | KLRC1,FCER1G,SYK |
| 0 | Immune system | Adaptive_NK | RAP1B | KLRC1,FCER1G,SYK |
| 0 | Immune system | Adaptive_NK | FLNA | KLRC1,FCER1G,SYK |
| 0 | Immune system | Adaptive_NK | EMP3 | KLRC1,FCER1G,SYK |
| 0 | Immune system | Adaptive_NK | B2M | KLRC1,FCER1G,SYK |
| 0 | Immune system | Adaptive_NK | C12orf75 | KLRC1,FCER1G,SYK |
| 0 | Immune system | Adaptive_NK | CALM1 | KLRC1,FCER1G,SYK |
| 0 | Immune system | Adaptive_NK | KLRD1 | KLRC1,FCER1G,SYK |
| 0 | Immune system | Adaptive_NK | GNG2 | KLRC1,FCER1G,SYK |
| 0 | Immune system | Adaptive_NK | TRBC1 | KLRC1,FCER1G,SYK |
| 0 | Immune system | Adaptive_NK | CCL4L2 | KLRC1,FCER1G,SYK |
| 0 | Immune system | Adaptive_NK | HLA-C | KLRC1,FCER1G,SYK |
| 0 | Immune system | Adaptive_NK | ACTG1 | KLRC1,FCER1G,SYK |
| 0 | Immune system | Adaptive_NK | ANXA1 | KLRC1,FCER1G,SYK |
| 0 | Immune system | Adaptive_NK | C1orf21 | KLRC1,FCER1G,SYK |
| 0 | Immune system | Adaptive_NK | SORL1 | KLRC1,FCER1G,SYK |
| 0 | Immune system | Adaptive_NK | HLA-E | KLRC1,FCER1G,SYK |
| 0 | Immune system | Adaptive_NK | AHNAK | KLRC1,FCER1G,SYK |
| 0 | Immune system | Adaptive_NK | HLA-B | KLRC1,FCER1G,SYK |
| 0 | Immune system | Adaptive_NK | ABHD17A | KLRC1,FCER1G,SYK |
| 0 | Immune system | Adaptive_NK | FGL2 | KLRC1,FCER1G,SYK |
| 0 | Immune system | Adaptive_NK | PPP2R5C | KLRC1,FCER1G,SYK |
| 0 | Immune system | Adaptive_NK | COL6A2 | KLRC1,FCER1G,SYK |

| | | | | |
|---|---|---|---|---|
| 0 | Immune system | Adaptive_NK | TSC22D3 | KLRC1,FCER1G,SYK |
| 0 | Immune system | Adaptive_NK | S100A6 | KLRC1,FCER1G,SYK |
| 0 | Immune system | Adaptive_NK | LYAR | KLRC1,FCER1G,SYK |
| 0 | Immune system | Adaptive_NK | GK5 | KLRC1,FCER1G,SYK |
| 0 | Immune system | Adaptive_NK | TPST2 | KLRC1,FCER1G,SYK |
| 0 | Immune system | Adaptive_NK | LPCAT1 | KLRC1,FCER1G,SYK |
| 0 | Immune system | Adaptive_NK | KIR2DL3 | KLRC1,FCER1G,SYK |
| 0 | Immune system | Adaptive_NK | CFL1 | KLRC1,FCER1G,SYK |
| 0 | Immune system | Adaptive_NK | SH3BP5 | KLRC1,FCER1G,SYK |
| 0 | Immune system | Adaptive_NK | XBP1 | KLRC1,FCER1G,SYK |
| 0 | Immune system | Adaptive_NK | ITGB2 | KLRC1,FCER1G,SYK |
| 0 | Immune system | Adaptive_NK | FCRL6 | KLRC1,FCER1G,SYK |
| 0 | Immune system | Adaptive_NK | PRKCB | KLRC1,FCER1G,SYK |
| 0 | Immune system | Adaptive_NK | MYL12A | KLRC1,FCER1G,SYK |
| 0 | Immune system | Adaptive_NK | HLA-DQA2 | KLRC1,FCER1G,SYK |
| 0 | Immune system | Adaptive_NK | MYBL1 | KLRC1,FCER1G,SYK |
| 0 | Immune system | Adaptive_NK | RAB29 | KLRC1,FCER1G,SYK |
| 0 | Immune system | Adaptive_NK | DSTN | KLRC1,FCER1G,SYK |
| 0 | Immune system | Adaptive_NK | PTGDR | KLRC1,FCER1G,SYK |
| 0 | Immune system | Adaptive_NK | RASGRP2 | KLRC1,FCER1G,SYK |
| 0 | Immune system | Adaptive_NK | MBP | KLRC1,FCER1G,SYK |
| 0 | Immune system | Adaptive_NK | F2R | KLRC1,FCER1G,SYK |
| 0 | Immune system | Adaptive_NK | ARPC2 | KLRC1,FCER1G,SYK |
| 0 | Immune system | Adaptive_NK | SYNE2 | KLRC1,FCER1G,SYK |
| 0 | Immune system | Adaptive_NK | LAIR1 | KLRC1,FCER1G,SYK |
| 0 | Immune system | Adaptive_NK | CCL4 | KLRC1,FCER1G,SYK |
| 0 | Immune system | Adaptive_NK | CD47 | KLRC1,FCER1G,SYK |
| 0 | Immune system | Adaptive_NK | TBX21 | KLRC1,FCER1G,SYK |
| 0 | Immune system | Adaptive_NK | CMC1 | KLRC1,FCER1G,SYK |
| 0 | Immune system | Adaptive_NK | ANXA4 | KLRC1,FCER1G,SYK |
| 0 | Immune system | Adaptive_NK | RGS9 | KLRC1,FCER1G,SYK |
| 0 | Immune system | Adaptive_NK | TMSB10 | KLRC1,FCER1G,SYK |
| 0 | Immune system | Adaptive_NK | PRDM1 | KLRC1,FCER1G,SYK |
| 0 | Immune system | Adaptive_NK | RASA3 | KLRC1,FCER1G,SYK |
| 0 | Immune system | Adaptive_NK | CD53 | KLRC1,FCER1G,SYK |
| 0 | Immune system | Adaptive_NK | SPN | KLRC1,FCER1G,SYK |
| 0 | Immune system | Adaptive_NK | ADRB2 | KLRC1,FCER1G,SYK |
| 0 | Immune system | Adaptive_NK | KLF3 | KLRC1,FCER1G,SYK |
| 0 | Immune system | Adaptive_NK | ADD3 | KLRC1,FCER1G,SYK |
| 0 | Immune system | Adaptive_NK | HLA-F | KLRC1,FCER1G,SYK |
| 0 | Immune system | Adaptive_NK | TFDP2 | KLRC1,FCER1G,SYK |
| 0 | Immune system | Adaptive_NK | SSBP3 | KLRC1,FCER1G,SYK |
| 0 | Immune system | Adaptive_NK | SYNE1 | KLRC1,FCER1G,SYK |
| 0 | Immune system | Adaptive_NK | SYTL3 | KLRC1,FCER1G,SYK |
| 0 | Immune system | Adaptive_NK | GLRX | KLRC1,FCER1G,SYK |
| 0 | Immune system | Adaptive_NK | HLA-DPB1 | KLRC1,FCER1G,SYK |
| 0 | Immune system | Adaptive_NK | RAP2B | KLRC1,FCER1G,SYK |
| 0 | Immune system | Adaptive_NK | UCP2 | KLRC1,FCER1G,SYK |
| 0 | Immune system | Adaptive_NK | TXNIP | KLRC1,FCER1G,SYK |
| 0 | Immune system | Adaptive_NK | VCL | KLRC1,FCER1G,SYK |
| 0 | Immune system | Adaptive_NK | MGAT4A | KLRC1,FCER1G,SYK |
| 0 | Immune system | Adaptive_NK | GZMA | KLRC1,FCER1G,SYK |
| 0 | Immune system | Adaptive_NK | GTF3C1 | KLRC1,FCER1G,SYK |
| 0 | Immune system | Adaptive_NK | PTGER2 | KLRC1,FCER1G,SYK |
| 0 | Immune system | Adaptive_NK | PTPRC | KLRC1,FCER1G,SYK |
| 0 | Immune system | Adaptive_NK | BIN2 | KLRC1,FCER1G,SYK |
| 0 | Immune system | Adaptive_NK | GIMAP4 | KLRC1,FCER1G,SYK |
| 0 | Immune system | Adaptive_NK | PXN | KLRC1,FCER1G,SYK |

| | | | | |
|---|---|---|---|---|
| 0 | Immune system | Adaptive_NK | ABI3 | KLRC1,FCER1G,SYK |
| 0 | Immune system | Adaptive_NK | ADGRE5 | KLRC1,FCER1G,SYK |
| 0 | Immune system | Adaptive_NK | SRGN | KLRC1,FCER1G,SYK |
| 0 | Immune system | Adaptive_NK | HDDC2 | KLRC1,FCER1G,SYK |
| 0 | Immune system | Adaptive_NK | ICAM2 | KLRC1,FCER1G,SYK |
| 0 | Immune system | Adaptive_NK | TPM4 | KLRC1,FCER1G,SYK |
| 0 | Immune system | Adaptive_NK | KLRC2 | KLRC1,FCER1G,SYK |
| 0 | Immune system | Adaptive_NK | CAP1 | KLRC1,FCER1G,SYK |
| 0 | Immune system | Adaptive_NK | LINC00861 | KLRC1,FCER1G,SYK |
| 0 | Immune system | Adaptive_NK | GRAP2 | KLRC1,FCER1G,SYK |
| 0 | Immune system | Adaptive_NK | EIF1 | KLRC1,FCER1G,SYK |
| 0 | Immune system | Adaptive_NK | IFNG | KLRC1,FCER1G,SYK |
| 0 | Immune system | Adaptive_NK | ITGAL | KLRC1,FCER1G,SYK |
| 0 | Immune system | Adaptive_NK | PTPRE | KLRC1,FCER1G,SYK |
| 0 | Immune system | Adaptive_NK | PTGER4 | KLRC1,FCER1G,SYK |
| 0 | Immune system | Adaptive_NK | HLA-DPA1 | KLRC1,FCER1G,SYK |
| 0 | Immune system | Adaptive_NK | CDC42SE1 | KLRC1,FCER1G,SYK |
| 0 | Immune system | Adaptive_NK | MAF | KLRC1,FCER1G,SYK |
| 0 | Immune system | Adaptive_NK | CTBP2 | KLRC1,FCER1G,SYK |
| 0 | Immune system | Adaptive_NK | CDKN2D | KLRC1,FCER1G,SYK |
| 0 | Immune system | Adaptive_NK | CYTH1 | KLRC1,FCER1G,SYK |
| 0 | Immune system | Adaptive_NK | MYO1G | KLRC1,FCER1G,SYK |
| 0 | Immune system | Adaptive_NK | SH3KBP1 | KLRC1,FCER1G,SYK |
| 0 | Immune system | Adaptive_NK | YWHAB | KLRC1,FCER1G,SYK |
| 0 | Immune system | Adaptive_NK | CDC25B | KLRC1,FCER1G,SYK |
| 0 | Immune system | Adaptive_NK | ATM | KLRC1,FCER1G,SYK |
| 0 | Immune system | Adaptive_NK | ACTR3 | KLRC1,FCER1G,SYK |
| 0 | Immune system | Adaptive_NK | RAB9A | KLRC1,FCER1G,SYK |
| 0 | Immune system | Adaptive_NK | RORA | KLRC1,FCER1G,SYK |
| 0 | Immune system | Adaptive_NK | GAB3 | KLRC1,FCER1G,SYK |
| 0 | Immune system | Adaptive_NK | KLF13 | KLRC1,FCER1G,SYK |
| 0 | Immune system | Adaptive_NK | KLF6 | KLRC1,FCER1G,SYK |
| 0 | Immune system | Adaptive_NK | CAST | KLRC1,FCER1G,SYK |
| 0 | Immune system | Adaptive_NK | IQGAP2 | KLRC1,FCER1G,SYK |
| 0 | Immune system | Adaptive_NK | RPL3 | KLRC1,FCER1G,SYK |
| 0 | Immune system | Adaptive_NK | CD320 | KLRC1,FCER1G,SYK |
| 0 | Immune system | Adaptive_NK | IGF2R | KLRC1,FCER1G,SYK |
| 0 | Immune system | Adaptive_NK | IFITM1 | KLRC1,FCER1G,SYK |
| 0 | Immune system | Adaptive_NK | GPSM3 | KLRC1,FCER1G,SYK |
| 0 | Immune system | Adaptive_NK | FTL | KLRC1,FCER1G,SYK |
| 0 | Immune system | Adaptive_NK | RHOG | KLRC1,FCER1G,SYK |
| 0 | Immune system | Adaptive_NK | CD3E | KLRC1,FCER1G,SYK |
| 0 | Immune system | Adaptive_NK | EBP | KLRC1,FCER1G,SYK |
| 0 | Immune system | Adaptive_NK | VASP | KLRC1,FCER1G,SYK |
| 0 | Immune system | Adaptive_NK | ARID5B | KLRC1,FCER1G,SYK |
| 0 | Immune system | Adaptive_NK | SH2D2A | KLRC1,FCER1G,SYK |
| 0 | Immune system | Adaptive_NK | MIAT | KLRC1,FCER1G,SYK |
| 0 | Immune system | Adaptive_NK | PTP4A2 | KLRC1,FCER1G,SYK |
| 0 | Immune system | Adaptive_NK | HSPA8 | KLRC1,FCER1G,SYK |
| 0 | Immune system | Adaptive_NK | IFITM2 | KLRC1,FCER1G,SYK |
| 0 | Immune system | Adaptive_NK | PYHIN1 | KLRC1,FCER1G,SYK |
| 0 | Immune system | Adaptive_NK | NDUFB2 | KLRC1,FCER1G,SYK |
| 0 | Immune system | Adaptive_NK | RPA2 | KLRC1,FCER1G,SYK |
| 0 | Immune system | Adaptive_NK | HIPK2 | KLRC1,FCER1G,SYK |
| 0 | Immune system | Adaptive_NK | SIGIRR | KLRC1,FCER1G,SYK |
| 0 | Immune system | Adaptive_NK | EIF4G3 | KLRC1,FCER1G,SYK |
| 0 | Immune system | Adaptive_NK | NDUFB7 | KLRC1,FCER1G,SYK |
| 0 | Immune system | Adaptive_NK | APMAP | KLRC1,FCER1G,SYK |

| | | | | |
|---|---|---|---|---|
| 0 | Immune system | Adaptive_NK | CD300A | KLRC1,FCER1G,SYK |
| 0 | Immune system | Adaptive_NK | CTSC | KLRC1,FCER1G,SYK |
| 0 | Immune system | Adaptive_NK | CCDC88C | KLRC1,FCER1G,SYK |
| 0 | Immune system | Adaptive_NK | PDIA3 | KLRC1,FCER1G,SYK |
| 0 | Immune system | Adaptive_NK | STK10 | KLRC1,FCER1G,SYK |
| 0 | Immune system | Adaptive_NK | TOB1 | KLRC1,FCER1G,SYK |
| 0 | Immune system | Adaptive_NK | YWHAZ | KLRC1,FCER1G,SYK |
| 0 | Immune system | Adaptive_NK | PPIB | KLRC1,FCER1G,SYK |
| 0 | Immune system | Adaptive_NK | METRNL | KLRC1,FCER1G,SYK |
| 0 | Immune system | Adaptive_NK | HLA-DRB1 | KLRC1,FCER1G,SYK |
| 0 | Immune system | Adaptive_NK | FAM49B | KLRC1,FCER1G,SYK |
| 0 | Immune system | Adaptive_NK | ARPC5 | KLRC1,FCER1G,SYK |
| 0 | Immune system | Adaptive_NK | MYADM | KLRC1,FCER1G,SYK |
| 0 | Immune system | Adaptive_NK | TERF1 | KLRC1,FCER1G,SYK |
| 0 | Immune system | Adaptive_NK | ARHGDIB | KLRC1,FCER1G,SYK |
| 0 | Immune system | Adaptive_NK | PPP1CA | KLRC1,FCER1G,SYK |
| 0 | Immune system | Adaptive_NK | PTPN12 | KLRC1,FCER1G,SYK |
| 0 | Immune system | Adaptive_NK | VAV3 | KLRC1,FCER1G,SYK |
| 0 | Immune system | Adaptive_NK | RTN4 | KLRC1,FCER1G,SYK |
| 0 | Immune system | Adaptive_NK | WDR1 | KLRC1,FCER1G,SYK |
| 0 | Immune system | Adaptive_NK | TLE4 | KLRC1,FCER1G,SYK |
| 0 | Immune system | Adaptive_NK | P4HB | KLRC1,FCER1G,SYK |
| 0 | Immune system | Adaptive_NK | ZBTB38 | KLRC1,FCER1G,SYK |
| 0 | Immune system | Adaptive_NK | CLIC3 | KLRC1,FCER1G,SYK |
| 0 | Immune system | Adaptive_NK | RGS19 | KLRC1,FCER1G,SYK |
| 0 | Immune system | Adaptive_NK | USP28 | KLRC1,FCER1G,SYK |
| 0 | Immune system | Adaptive_NK | FGR | KLRC1,FCER1G,SYK |
| 0 | Immune system | Adaptive_NK | TNFRSF1B | KLRC1,FCER1G,SYK |
| 0 | Immune system | Adaptive_NK | RPL21 | KLRC1,FCER1G,SYK |
| 0 | Immune system | Adaptive_NK | OSTF1 | KLRC1,FCER1G,SYK |
| 0 | Immune system | Adaptive_NK | PRR5L | KLRC1,FCER1G,SYK |
| 0 | Immune system | Adaptive_NK | HSPA5 | KLRC1,FCER1G,SYK |
| 0 | Immune system | Adaptive_NK | TMBIM6 | KLRC1,FCER1G,SYK |
| 0 | Immune system | Adaptive_NK | DBI | KLRC1,FCER1G,SYK |
| 0 | Immune system | Adaptive_NK | TMEM181 | KLRC1,FCER1G,SYK |
| 0 | Immune system | Adaptive_NK | ACTR2 | KLRC1,FCER1G,SYK |
| 0 | Immune system | Adaptive_NK | CORO1A | KLRC1,FCER1G,SYK |
| 0 | Immune system | Adaptive_NK | RAC2 | KLRC1,FCER1G,SYK |
| 0 | Immune system | Adaptive_NK | ARHGAP25 | KLRC1,FCER1G,SYK |
| 0 | Immune system | Adaptive_NK | UPP1 | KLRC1,FCER1G,SYK |
| 0 | Immune system | Adaptive_NK | LYN | KLRC1,FCER1G,SYK |
| 0 | Immune system | Adaptive_NK | TMEM173 | KLRC1,FCER1G,SYK |
| 0 | Immune system | Adaptive_NK | AES | KLRC1,FCER1G,SYK |
| 0 | Immune system | Adaptive_NK | MYO1F | KLRC1,FCER1G,SYK |
| 0 | Immune system | Adaptive_NK | SASH3 | KLRC1,FCER1G,SYK |
| 0 | Immune system | Adaptive_NK | BATF | KLRC1,FCER1G,SYK |
| 0 | Immune system | Adaptive_NK | GIMAP1 | KLRC1,FCER1G,SYK |
| 0 | Immune system | Adaptive_NK | UBE2F | KLRC1,FCER1G,SYK |
| 0 | Immune system | Adaptive_NK | CAPN2 | KLRC1,FCER1G,SYK |
| 0 | Immune system | Adaptive_NK | SLC9A3R1 | KLRC1,FCER1G,SYK |
| 0 | Immune system | Adaptive_NK | SPCS3 | KLRC1,FCER1G,SYK |
| 0 | Immune system | Adaptive_NK | GMFG | KLRC1,FCER1G,SYK |
| 0 | Immune system | Adaptive_NK | SMAP2 | KLRC1,FCER1G,SYK |
| 0 | Immune system | Adaptive_NK | HNRNPF | KLRC1,FCER1G,SYK |
| 0 | Immune system | Adaptive_NK | CD99 | KLRC1,FCER1G,SYK |
| 0 | Immune system | Adaptive_NK | CDC42 | KLRC1,FCER1G,SYK |
| 0 | Immune system | Adaptive_NK | RHOA | KLRC1,FCER1G,SYK |
| 0 | Immune system | Adaptive_NK | ARPC4 | KLRC1,FCER1G,SYK |

| | | | | |
|---|---|---|---|---|
| 0 | Immune system | Adaptive_NK | BIN1 | KLRC1,FCER1G,SYK |
| 0 | Immune system | Adaptive_NK | ORAI1 | KLRC1,FCER1G,SYK |
| 0 | Immune system | Adaptive_NK | DHRS7 | KLRC1,FCER1G,SYK |
| 0 | Immune system | Adaptive_NK | ANXA2 | KLRC1,FCER1G,SYK |
| 0 | Immune system | Adaptive_NK | TPP1 | KLRC1,FCER1G,SYK |
| 0 | Immune system | Adaptive_NK | ATP2B4 | KLRC1,FCER1G,SYK |
| 0 | Immune system | Adaptive_NK | CDC42EP3 | KLRC1,FCER1G,SYK |
| 0 | Immune system | Adaptive_NK | UTRN | KLRC1,FCER1G,SYK |
| 0 | Immune system | Adaptive_NK | GNAI2 | KLRC1,FCER1G,SYK |
| 0 | Immune system | Adaptive_NK | FYN | KLRC1,FCER1G,SYK |
| 0 | Immune system | Adaptive_NK | TES | KLRC1,FCER1G,SYK |
| 0 | Immune system | Adaptive_NK | SDF2L1 | KLRC1,FCER1G,SYK |
| 0 | Immune system | Adaptive_NK | TGFBR1 | KLRC1,FCER1G,SYK |
| 0 | Immune system | Adaptive_NK | RNF19A | KLRC1,FCER1G,SYK |
| 0 | Immune system | Adaptive_NK | KLRG1 | KLRC1,FCER1G,SYK |
| 0 | Immune system | Adaptive_NK | PARP15 | KLRC1,FCER1G,SYK |
| 0 | Immune system | Adaptive_NK | MED15 | KLRC1,FCER1G,SYK |
| 0 | Immune system | Adaptive_NK | SLC15A4 | KLRC1,FCER1G,SYK |
| 0 | Immune system | Adaptive_NK | S1PR4 | KLRC1,FCER1G,SYK |
| 0 | Immune system | Adaptive_NK | LLGL2 | KLRC1,FCER1G,SYK |
| 0 | Immune system | Adaptive_NK | CALR | KLRC1,FCER1G,SYK |
| 0 | Immune system | Adaptive_NK | SFT2D1 | KLRC1,FCER1G,SYK |
| 0 | Immune system | Adaptive_NK | MIEN1 | KLRC1,FCER1G,SYK |
| 0 | Immune system | Adaptive_NK | TUBA4A | KLRC1,FCER1G,SYK |
| 0 | Immune system | Adaptive_NK | NCOA1 | KLRC1,FCER1G,SYK |
| 0 | Immune system | Adaptive_NK | SUN2 | KLRC1,FCER1G,SYK |
| 0 | Immune system | Adaptive_NK | UHMK1 | KLRC1,FCER1G,SYK |
| 0 | Immune system | Adaptive_NK | ANXA6 | KLRC1,FCER1G,SYK |
| 0 | Immune system | Adaptive_NK | MIDN | KLRC1,FCER1G,SYK |
| 0 | Immune system | Adaptive_NK | TNFRSF14 | KLRC1,FCER1G,SYK |
| 0 | Immune system | Adaptive_NK | PTPN18 | KLRC1,FCER1G,SYK |
| 0 | Immune system | Adaptive_NK | GPR65 | KLRC1,FCER1G,SYK |
| 0 | Immune system | Adaptive_NK | AKNA | KLRC1,FCER1G,SYK |
| 0 | Immune system | Adaptive_NK | MANF | KLRC1,FCER1G,SYK |
| 0 | Immune system | Adaptive_NK | LCK | KLRC1,FCER1G,SYK |
| 0 | Immune system | Adaptive_NK | LMAN2 | KLRC1,FCER1G,SYK |
| 0 | Immune system | Adaptive_NK | MOB3A | KLRC1,FCER1G,SYK |
| 0 | Immune system | Adaptive_NK | PRELID1 | KLRC1,FCER1G,SYK |
| 0 | Immune system | Adaptive_NK | CAPZB | KLRC1,FCER1G,SYK |
| 0 | Immune system | Adaptive_NK | CREM | KLRC1,FCER1G,SYK |
| 0 | Immune system | Adaptive_NK | FBXW5 | KLRC1,FCER1G,SYK |
| 0 | Immune system | Adaptive_NK | CDK2AP2 | KLRC1,FCER1G,SYK |
| 0 | Immune system | Adaptive_NK | SH3BP2 | KLRC1,FCER1G,SYK |
| 0 | Immune system | Adaptive_NK | MSN | KLRC1,FCER1G,SYK |
| 0 | Immune system | Adaptive_NK | PPIA | KLRC1,FCER1G,SYK |
| 0 | Immune system | Adaptive_NK | YWHAQ | KLRC1,FCER1G,SYK |
| 0 | Immune system | Adaptive_NK | PRMT2 | KLRC1,FCER1G,SYK |
| 0 | Immune system | Adaptive_NK | ATP1A1 | KLRC1,FCER1G,SYK |
| 0 | Immune system | Adaptive_NK | CLIC1 | KLRC1,FCER1G,SYK |
| 0 | Immune system | Adaptive_NK | STK38 | KLRC1,FCER1G,SYK |
| 0 | Immune system | Adaptive_NK | DIAPH1 | KLRC1,FCER1G,SYK |
| 0 | Immune system | Adaptive_NK | DENND2D | KLRC1,FCER1G,SYK |
| 0 | Immune system | Adaptive_NK | S100A10 | KLRC1,FCER1G,SYK |
| 0 | Immune system | Adaptive_NK | SUPT4H1 | KLRC1,FCER1G,SYK |
| 0 | Immune system | Adaptive_NK | ATP2B1-AS1 | KLRC1,FCER1G,SYK |
| 0 | Immune system | Adaptive_NK | SYNGR1 | KLRC1,FCER1G,SYK |
| 0 | Immune system | Adaptive_NK | MAPRE2 | KLRC1,FCER1G,SYK |
| 0 | Immune system | Adaptive_NK | HSP90B1 | KLRC1,FCER1G,SYK |

| | | | | |
|---|---|---|---|---|
| 0 | Immune system | Adaptive_NK | G6PD | KLRC1,FCER1G,SYK |
| 0 | Immune system | Adaptive_NK | TBCB | KLRC1,FCER1G,SYK |
| 0 | Immune system | Adaptive_NK | TRPV2 | KLRC1,FCER1G,SYK |
| 0 | Immune system | Adaptive_NK | GRK6 | KLRC1,FCER1G,SYK |
| 0 | Immune system | Adaptive_NK | TIMP1 | KLRC1,FCER1G,SYK |
| 0 | Immune system | Adaptive_NK | MTPN | KLRC1,FCER1G,SYK |
| 0 | Immune system | Adaptive_NK | LDLR | KLRC1,FCER1G,SYK |
| 0 | Immune system | Adaptive_NK | ZAP70 | KLRC1,FCER1G,SYK |
| 0 | Immune system | Adaptive_NK | C12orf57 | KLRC1,FCER1G,SYK |
| 0 | Immune system | Adaptive_NK | CISD3 | KLRC1,FCER1G,SYK |
| 0 | Immune system | Adaptive_NK | RNF167 | KLRC1,FCER1G,SYK |
| 0 | Immune system | Adaptive_NK | IVNS1ABP | KLRC1,FCER1G,SYK |
| 0 | Immune system | Adaptive_NK | ARL6IP1 | KLRC1,FCER1G,SYK |
| 0 | Immune system | Adaptive_NK | PPP1R18 | KLRC1,FCER1G,SYK |
| 0 | Immune system | Adaptive_NK | TPM3 | KLRC1,FCER1G,SYK |
| 0 | Immune system | Adaptive_NK | GIMAP7 | KLRC1,FCER1G,SYK |
| 0 | Immune system | Adaptive_NK | SNF8 | KLRC1,FCER1G,SYK |
| 0 | Immune system | Adaptive_NK | SEC11C | KLRC1,FCER1G,SYK |
| 0 | Immune system | Adaptive_NK | LBR | KLRC1,FCER1G,SYK |
| 0 | Immune system | Adaptive_NK | RAB8A | KLRC1,FCER1G,SYK |
| 0 | Immune system | Adaptive_NK | ZNF276 | KLRC1,FCER1G,SYK |
| 0 | Immune system | Adaptive_NK | PIP4K2A | KLRC1,FCER1G,SYK |
| 0 | Immune system | Adaptive_NK | RPS26 | KLRC1,FCER1G,SYK |
| 0 | Immune system | Adaptive_NK | RASSF1 | KLRC1,FCER1G,SYK |
| 0 | Immune system | Adaptive_NK | BCL11B | KLRC1,FCER1G,SYK |
| 0 | Immune system | Adaptive_NK | PCBP1 | KLRC1,FCER1G,SYK |
| 0 | Immune system | Adaptive_NK | FKBP11 | KLRC1,FCER1G,SYK |
| 0 | Immune system | Adaptive_NK | TMEM50A | KLRC1,FCER1G,SYK |
| 0 | Immune system | Adaptive_NK | MYDGF | KLRC1,FCER1G,SYK |
| 0 | Immune system | Adaptive_NK | RNF126 | KLRC1,FCER1G,SYK |
| 0 | Immune system | Adaptive_NK | RRBP1 | KLRC1,FCER1G,SYK |
| 0 | Immune system | Adaptive_NK | PLEKHA1 | KLRC1,FCER1G,SYK |
| 0 | Immune system | Adaptive_NK | SELPLG | KLRC1,FCER1G,SYK |
| 0 | Immune system | Adaptive_NK | SH2D1B | KLRC1,FCER1G,SYK |
| 0 | Immune system | Adaptive_NK | RAB5C | KLRC1,FCER1G,SYK |
| 0 | Immune system | Adaptive_NK | ARRDC3 | KLRC1,FCER1G,SYK |
| 0 | Immune system | Adaptive_NK | ARL6IP5 | KLRC1,FCER1G,SYK |
| 0 | Immune system | Adaptive_NK | TMEM2 | KLRC1,FCER1G,SYK |
| 0 | Immune system | Adaptive_NK | CCDC85B | KLRC1,FCER1G,SYK |
| 0 | Immune system | Adaptive_NK | ARPC1B | KLRC1,FCER1G,SYK |
| 0 | Immune system | Adaptive_NK | SIRT2 | KLRC1,FCER1G,SYK |
| 0 | Immune system | Adaptive_NK | CD226 | KLRC1,FCER1G,SYK |
| 0 | Immune system | Adaptive_NK | LRRFIP1 | KLRC1,FCER1G,SYK |
| 0 | Immune system | Adaptive_NK | RAB10 | KLRC1,FCER1G,SYK |
| 0 | Immune system | Adaptive_NK | ATP1B3 | KLRC1,FCER1G,SYK |
| 0 | Immune system | Adaptive_NK | PDIA6 | KLRC1,FCER1G,SYK |
| 0 | Immune system | Adaptive_NK | SRPK2 | KLRC1,FCER1G,SYK |
| 0 | Immune system | Adaptive_NK | IQGAP1 | KLRC1,FCER1G,SYK |
| 0 | Immune system | Adaptive_NK | PGAM1 | KLRC1,FCER1G,SYK |
| 0 | Immune system | Adaptive_NK | CAPNS1 | KLRC1,FCER1G,SYK |
| 0 | Immune system | Adaptive_NK | TMED9 | KLRC1,FCER1G,SYK |
| 0 | Immune system | Adaptive_NK | MFSD10 | KLRC1,FCER1G,SYK |
| 0 | Immune system | Adaptive_NK | ATP5F1 | KLRC1,FCER1G,SYK |
| 0 | Immune system | Adaptive_NK | DECR1 | KLRC1,FCER1G,SYK |
| 0 | Immune system | Adaptive_NK | PDLIM2 | KLRC1,FCER1G,SYK |
| 0 | Immune system | Adaptive_NK | SERPINB1 | KLRC1,FCER1G,SYK |
| 0 | Immune system | Adaptive_NK | CCDC82 | KLRC1,FCER1G,SYK |
| 0 | Immune system | Adaptive_NK | CEBPB | KLRC1,FCER1G,SYK |

| | | | | |
|---|---|---|---|---|
| 0 | Immune system | Adaptive_NK | ITGB1BP1 | KLRC1,FCER1G,SYK |
| 0 | Immune system | Adaptive_NK | M6PR | KLRC1,FCER1G,SYK |
| 0 | Immune system | Adaptive_NK | TMEM59 | KLRC1,FCER1G,SYK |
| 0 | Immune system | Adaptive_NK | RECQL | KLRC1,FCER1G,SYK |
| 0 | Immune system | Adaptive_NK | RGS14 | KLRC1,FCER1G,SYK |
| 0 | Immune system | Adaptive_NK | PLEKHF1 | KLRC1,FCER1G,SYK |
| 0 | Immune system | Adaptive_NK | NFATC2 | KLRC1,FCER1G,SYK |
| 0 | Immune system | Adaptive_NK | PPP2R5A | KLRC1,FCER1G,SYK |
| 0 | Immune system | Adaptive_NK | SEC61B | KLRC1,FCER1G,SYK |
| 0 | Immune system | Adaptive_NK | SYTL1 | KLRC1,FCER1G,SYK |
| 0 | Immune system | Adaptive_NK | SUB1 | KLRC1,FCER1G,SYK |
| 0 | Immune system | Adaptive_NK | TLN1 | KLRC1,FCER1G,SYK |
| 0 | Immune system | Adaptive_NK | SPCS2 | KLRC1,FCER1G,SYK |
| 0 | Immune system | Adaptive_NK | PDAP1 | KLRC1,FCER1G,SYK |
| 0 | Immune system | Adaptive_NK | VAMP2 | KLRC1,FCER1G,SYK |
| 0 | Immune system | Adaptive_NK | SLAMF7 | KLRC1,FCER1G,SYK |
| 0 | Immune system | Adaptive_NK | CRBN | KLRC1,FCER1G,SYK |
| 0 | Immune system | Adaptive_NK | SELENOT | KLRC1,FCER1G,SYK |
| 0 | Immune system | Adaptive_NK | TECR | KLRC1,FCER1G,SYK |
| 0 | Immune system | Adaptive_NK | RNF149 | KLRC1,FCER1G,SYK |
| 0 | Immune system | Adaptive_NK | SIPA1 | KLRC1,FCER1G,SYK |
| 0 | Immune system | Adaptive_NK | EIF4EBP2 | KLRC1,FCER1G,SYK |
| 0 | Immune system | Adaptive_NK | HSH2D | KLRC1,FCER1G,SYK |
| 0 | Immune system | Adaptive_NK | IMP3 | KLRC1,FCER1G,SYK |
| 0 | Immune system | Adaptive_NK | CIB1 | KLRC1,FCER1G,SYK |
| 0 | Immune system | Adaptive_NK | LRP10 | KLRC1,FCER1G,SYK |
| 0 | Immune system | Adaptive_NK | ARPC5L | KLRC1,FCER1G,SYK |
| 0 | Immune system | Adaptive_NK | RAPGEF1 | KLRC1,FCER1G,SYK |
| 0 | Immune system | Adaptive_NK | POLR3GL | KLRC1,FCER1G,SYK |
| 0 | Immune system | Adaptive_NK | RNF169 | KLRC1,FCER1G,SYK |
| 0 | Immune system | Adaptive_NK | ZFAND6 | KLRC1,FCER1G,SYK |
| 0 | Immune system | Adaptive_NK | ARHGAP30 | KLRC1,FCER1G,SYK |
| 0 | Immune system | Adaptive_NK | STOM | KLRC1,FCER1G,SYK |
| 0 | Immune system | Adaptive_NK | TCF25 | KLRC1,FCER1G,SYK |
| 0 | Immune system | Adaptive_NK | CTDSP1 | KLRC1,FCER1G,SYK |
| 0 | Immune system | Adaptive_NK | PSMB10 | KLRC1,FCER1G,SYK |
| 0 | Immune system | Adaptive_NK | ATOX1 | KLRC1,FCER1G,SYK |
| 0 | Immune system | Adaptive_NK | SYAP1 | KLRC1,FCER1G,SYK |
| 0 | Immune system | Adaptive_NK | CYB561D2 | KLRC1,FCER1G,SYK |
| 0 | Immune system | Adaptive_NK | EIF4B | KLRC1,FCER1G,SYK |
| 0 | Immune system | Adaptive_NK | TAF7 | KLRC1,FCER1G,SYK |
| 0 | Immune system | Adaptive_NK | SRPRA | KLRC1,FCER1G,SYK |
| 0 | Immune system | Adaptive_NK | CANX | KLRC1,FCER1G,SYK |
| 0 | Immune system | Adaptive_NK | RNASEH2C | KLRC1,FCER1G,SYK |
| 0 | Immune system | Adaptive_NK | IDI1 | KLRC1,FCER1G,SYK |
| 0 | Immune system | Adaptive_NK | SSBP4 | KLRC1,FCER1G,SYK |
| 0 | Immune system | Adaptive_NK | C16orf54 | KLRC1,FCER1G,SYK |
| 0 | Immune system | Adaptive_NK | CYFIP2 | KLRC1,FCER1G,SYK |
| 0 | Immune system | Adaptive_NK | RNPEPL1 | KLRC1,FCER1G,SYK |
| 0 | Immune system | Adaptive_NK | ARHGDIA | KLRC1,FCER1G,SYK |
| 0 | Immune system | Adaptive_NK | CELF2 | KLRC1,FCER1G,SYK |
| 0 | Immune system | Adaptive_NK | PMAIP1 | KLRC1,FCER1G,SYK |
| 0 | Immune system | Adaptive_NK | FKBP2 | KLRC1,FCER1G,SYK |
| 0 | Immune system | Adaptive_NK | BANF1 | KLRC1,FCER1G,SYK |
| 0 | Immune system | Adaptive_NK | PAXX | KLRC1,FCER1G,SYK |
| 0 | Immune system | Adaptive_NK | TAOK3 | KLRC1,FCER1G,SYK |
| 0 | Immune system | Adaptive_NK | CFLAR | KLRC1,FCER1G,SYK |
| 0 | Immune system | Adaptive_NK | KRT10 | KLRC1,FCER1G,SYK |

| | | | | |
|---|---|---|---|---|
| 0 | Immune system | Adaptive_NK | SELENOF | KLRC1,FCER1G,SYK |
| 0 | Immune system | Adaptive_NK | HIPK1 | KLRC1,FCER1G,SYK |
| 0 | Immune system | Adaptive_NK | BAZ1A | KLRC1,FCER1G,SYK |
| 0 | Immune system | Adaptive_NK | UBE2Q1 | KLRC1,FCER1G,SYK |
| 0 | Immune system | Adaptive_NK | TRAPPC10 | KLRC1,FCER1G,SYK |
| 0 | Immune system | Adaptive_NK | MORC3 | KLRC1,FCER1G,SYK |
| 0 | Immune system | Adaptive_NK | AKAP13 | KLRC1,FCER1G,SYK |
| 0 | Immune system | Adaptive_NK | PHF20 | KLRC1,FCER1G,SYK |
| 0 | Immune system | Adaptive_NK | EIF3K | KLRC1,FCER1G,SYK |
| 0 | Immune system | Adaptive_NK | WASF2 | KLRC1,FCER1G,SYK |
| 0 | Immune system | Adaptive_NK | ERBIN | KLRC1,FCER1G,SYK |
| 0 | Immune system | Adaptive_NK | BZW1 | KLRC1,FCER1G,SYK |
| 0 | Immune system | Adaptive_NK | LAT | KLRC1,FCER1G,SYK |
| 0 | Immune system | Adaptive_NK | SERPINB6 | KLRC1,FCER1G,SYK |
| 0 | Immune system | Adaptive_NK | DOCK11 | KLRC1,FCER1G,SYK |
| 0 | Immune system | Adaptive_NK | C1QBP | KLRC1,FCER1G,SYK |
| 0 | Immune system | Adaptive_NK | TROVE2 | KLRC1,FCER1G,SYK |
| 0 | Immune system | Adaptive_NK | LINC00869 | KLRC1,FCER1G,SYK |
| 0 | Immune system | Adaptive_NK | SELENOW | KLRC1,FCER1G,SYK |
| 0 | Immune system | Adaptive_NK | ARRB2 | KLRC1,FCER1G,SYK |
| 0 | Immune system | Adaptive_NK | TINF2 | KLRC1,FCER1G,SYK |
| 0 | Immune system | Adaptive_NK | MIB2 | KLRC1,FCER1G,SYK |
| 0 | Immune system | Adaptive_NK | TCP1 | KLRC1,FCER1G,SYK |
| 0 | Immune system | Adaptive_NK | CMPK1 | KLRC1,FCER1G,SYK |
| 0 | Immune system | Adaptive_NK | RNF125 | KLRC1,FCER1G,SYK |
| 0 | Immune system | Adaptive_NK | ZBTB7A | KLRC1,FCER1G,SYK |
| 0 | Immune system | Adaptive_NK | PTPN4 | KLRC1,FCER1G,SYK |
| 0 | Immune system | Adaptive_NK | PRNP | KLRC1,FCER1G,SYK |
| 0 | Immune system | Adaptive_NK | TADA3 | KLRC1,FCER1G,SYK |
| 0 | Immune system | Adaptive_NK | FERMT3 | KLRC1,FCER1G,SYK |
| 0 | Immune system | Adaptive_NK | NCR3 | KLRC1,FCER1G,SYK |
| 0 | Immune system | Adaptive_NK | TMED5 | KLRC1,FCER1G,SYK |
| 0 | Immune system | Adaptive_NK | GLG1 | KLRC1,FCER1G,SYK |
| 0 | Immune system | Adaptive_NK | TMEM9B | KLRC1,FCER1G,SYK |
| 0 | Immune system | Adaptive_NK | COX8A | KLRC1,FCER1G,SYK |
| 0 | Immune system | Adaptive_NK | RAC1 | KLRC1,FCER1G,SYK |
| 0 | Immune system | Adaptive_NK | SRSF9 | KLRC1,FCER1G,SYK |
| 0 | Immune system | Adaptive_NK | TTC16 | KLRC1,FCER1G,SYK |
| 0 | Immune system | Adaptive_NK | CADM1 | KLRC1,FCER1G,SYK |
| 0 | Immune system | Adaptive_NK | SGCD | KLRC1,FCER1G,SYK |
| 0 | Immune system | Adaptive_NK | DCP1B | KLRC1,FCER1G,SYK |
| 0 | Immune system | Adaptive_NK | DRAXIN | KLRC1,FCER1G,SYK |
| 0 | Immune system | Adaptive_NK | CAMK2N1 | KLRC1,FCER1G,SYK |
| 0 | Immune system | Adaptive_NK | LOC283177 | KLRC1,FCER1G,SYK |
| 0 | Immune system | Adaptive_NK | NCAPH | KLRC1,FCER1G,SYK |
| 0 | Immune system | Adaptive_NK | MYO6 | KLRC1,FCER1G,SYK |
| 0 | Immune system | Adaptive_NK | SBK1 | KLRC1,FCER1G,SYK |
| 0 | Immune system | Adaptive_NK | CCL5 | KLRC1,FCER1G,SYK |
| 0 | Immune system | Adaptive_NK | RAB11FIP5 | KLRC1,FCER1G,SYK |
| 0 | Immune system | Adaptive_NK | JAKMIP1 | KLRC1,FCER1G,SYK |
| 0 | Immune system | Adaptive_NK | CORO2A | KLRC1,FCER1G,SYK |
| 0 | Immune system | Adaptive_NK | B3GAT1 | KLRC1,FCER1G,SYK |
| 0 | Immune system | Adaptive_NK | SATB2 | KLRC1,FCER1G,SYK |
| 0 | Immune system | Adaptive_NK | PDGFRB | KLRC1,FCER1G,SYK |
| 0 | Immune system | Adaptive_NK | CD6 | KLRC1,FCER1G,SYK |
| 0 | Immune system | Adaptive_NK | EPB41L4A | KLRC1,FCER1G,SYK |
| 0 | Immune system | Adaptive_NK | GDPD5 | KLRC1,FCER1G,SYK |
| 0 | Immune system | Adaptive_NK | F8 | KLRC1,FCER1G,SYK |

| | | | | |
|---|---|---|---|---|
| 0 | Immune system | Adaptive_NK | LMTK3 | KLRC1,FCER1G,SYK |
| 0 | Immune system | Adaptive_NK | RCAN2 | KLRC1,FCER1G,SYK |
| 0 | Immune system | Adaptive_NK | GOLM1 | KLRC1,FCER1G,SYK |
| 0 | Immune system | Adaptive_NK | ITPRIPL1 | KLRC1,FCER1G,SYK |
| 0 | Immune system | Adaptive_NK | NUAK1 | KLRC1,FCER1G,SYK |
| 0 | Immune system | Adaptive_NK | KLRAP1 | KLRC1,FCER1G,SYK |
| 0 | Immune system | Adaptive_NK | TRG-AS1 | KLRC1,FCER1G,SYK |
| 0 | Immune system | Adaptive_NK | PPFIA3 | KLRC1,FCER1G,SYK |
| 0 | Immune system | Adaptive_NK | WNT10B | KLRC1,FCER1G,SYK |
| 0 | Immune system | Adaptive_NK | TPRG1 | KLRC1,FCER1G,SYK |
| 0 | Immune system | Adaptive_NK | CCDC85C | KLRC1,FCER1G,SYK |
| 0 | Immune system | Adaptive_NK | LINC00944 | KLRC1,FCER1G,SYK |
| 0 | Immune system | Adaptive_NK | NSG1 | KLRC1,FCER1G,SYK |
| 0 | Immune system | Adaptive_NK | CDKN2A | KLRC1,FCER1G,SYK |
| 0 | Immune system | Adaptive_NK | DUSP8 | KLRC1,FCER1G,SYK |
| 0 | Immune system | Adaptive_NK | ACTA2 | KLRC1,FCER1G,SYK |
| 0 | Immune system | Adaptive_NK | CD2 | KLRC1,FCER1G,SYK |
| 0 | Immune system | Adaptive_NK | PTMS | KLRC1,FCER1G,SYK |
| 0 | Immune system | Adaptive_NK | ABCD2 | KLRC1,FCER1G,SYK |
| 0 | Immune system | Adaptive_NK | TP53TG1 | KLRC1,FCER1G,SYK |
| 0 | Immune system | Adaptive_NK | LRRC16B | KLRC1,FCER1G,SYK |
| 0 | Immune system | Adaptive_NK | KIAA1671 | KLRC1,FCER1G,SYK |
| 0 | Immune system | Adaptive_NK | PAK6 | KLRC1,FCER1G,SYK |
| 0 | Immune system | Adaptive_NK | MXRA7 | KLRC1,FCER1G,SYK |
| 0 | Immune system | Adaptive_NK | FAM131B | KLRC1,FCER1G,SYK |
| 0 | Immune system | Adaptive_NK | ATP8B2 | KLRC1,FCER1G,SYK |
| 0 | Immune system | Adaptive_NK | FAM53B | KLRC1,FCER1G,SYK |
| 0 | Immune system | Adaptive_NK | GLB1L2 | KLRC1,FCER1G,SYK |
| 0 | Immune system | Adaptive_NK | CDC14B | KLRC1,FCER1G,SYK |
| 0 | Immune system | Adaptive_NK | PCNXL2 | KLRC1,FCER1G,SYK |
| 0 | Immune system | Adaptive_NK | HOXC5 | KLRC1,FCER1G,SYK |
| 0 | Immune system | Adaptive_NK | MCOLN2 | KLRC1,FCER1G,SYK |
| 0 | Immune system | Adaptive_NK | MVB12B | KLRC1,FCER1G,SYK |
| 0 | Immune system | Adaptive_NK | ZNF365 | KLRC1,FCER1G,SYK |
| 0 | Immune system | Adaptive_NK | SOX13 | KLRC1,FCER1G,SYK |
| 0 | Immune system | Adaptive_NK | GCNT4 | KLRC1,FCER1G,SYK |
| 0 | Immune system | Adaptive_NK | KLRC3 | KLRC1,FCER1G,SYK |
| 0 | Immune system | Adaptive_NK | LAG3 | KLRC1,FCER1G,SYK |
| 0 | Immune system | Adaptive_NK | TMEM255A | KLRC1,FCER1G,SYK |
| 0 | Immune system | Adaptive_NK | GOLGA7B | KLRC1,FCER1G,SYK |
| 0 | Immune system | Adaptive_NK | NINL | KLRC1,FCER1G,SYK |
| 0 | Immune system | Adaptive_NK | DAPK2 | KLRC1,FCER1G,SYK |
| 0 | Immune system | Adaptive_NK | ARHGEF28 | KLRC1,FCER1G,SYK |
| 0 | Immune system | Adaptive_NK | GNAO1 | KLRC1,FCER1G,SYK |
| 0 | Immune system | Adaptive_NK | PBX4 | KLRC1,FCER1G,SYK |
| 0 | Immune system | Adaptive_NK | CRIP1 | KLRC1,FCER1G,SYK |
| 0 | Immune system | Adaptive_NK | EPB41L4A-AS2 | KLRC1,FCER1G,SYK |
| 0 | Immune system | Adaptive_NK | LINC00943 | KLRC1,FCER1G,SYK |
| 0 | Immune system | Adaptive_NK | PATL2 | KLRC1,FCER1G,SYK |
| 0 | Immune system | Adaptive_NK | FKBP1B | KLRC1,FCER1G,SYK |
| 0 | Immune system | Adaptive_NK | MLF1 | KLRC1,FCER1G,SYK |
| 0 | Immune system | Adaptive_NK | EPN2 | KLRC1,FCER1G,SYK |
| 0 | Immune system | Adaptive_NK | KIF5A | KLRC1,FCER1G,SYK |
| 0 | Immune system | Adaptive_NK | KCNA3 | KLRC1,FCER1G,SYK |
| 0 | Immune system | Adaptive_NK | LRFN2 | KLRC1,FCER1G,SYK |
| 0 | Immune system | Adaptive_NK | ISL2 | KLRC1,FCER1G,SYK |
| 0 | Immune system | Adaptive_NK | LIME1 | KLRC1,FCER1G,SYK |
| 0 | Immune system | Adaptive_NK | GPR153 | KLRC1,FCER1G,SYK |

| | | | | |
|---|---|---|---|---|
| 0 | Immune system | Adaptive_NK | KLHL4 | KLRC1,FCER1G,SYK |
| 0 | Immune system | Adaptive_NK | KIAA1324 | KLRC1,FCER1G,SYK |
| 0 | Immune system | Adaptive_NK | VSTM2B | KLRC1,FCER1G,SYK |
| 0 | Immune system | Adaptive_NK | CDYL2 | KLRC1,FCER1G,SYK |
| 0 | Immune system | Adaptive_NK | SLC35G2 | KLRC1,FCER1G,SYK |
| 0 | Immune system | Adaptive_NK | ATL1 | KLRC1,FCER1G,SYK |
| 0 | Immune system | Adaptive_NK | FRMPD3 | KLRC1,FCER1G,SYK |
| 0 | Immune system | Adaptive_NK | CXXC4 | KLRC1,FCER1G,SYK |
| 0 | Immune system | Adaptive_NK | CLTCL1 | KLRC1,FCER1G,SYK |
| 0 | Immune system | Adaptive_NK | FAM167A | KLRC1,FCER1G,SYK |
| 0 | Immune system | Adaptive_NK | PPP2R2B | KLRC1,FCER1G,SYK |
| 0 | Immune system | Adaptive_NK | LRFN3 | KLRC1,FCER1G,SYK |
| 0 | Immune system | Adaptive_NK | HOXC4 | KLRC1,FCER1G,SYK |
| 0 | Immune system | Adaptive_NK | TCERG1L | KLRC1,FCER1G,SYK |
| 0 | Immune system | Adaptive_NK | C1orf177 | KLRC1,FCER1G,SYK |
| 0 | Immune system | Adaptive_NK | TSHZ3 | KLRC1,FCER1G,SYK |
| 0 | Immune system | Adaptive_NK | FOXD1 | KLRC1,FCER1G,SYK |
| 0 | Immune system | Adaptive_NK | MYO3B | KLRC1,FCER1G,SYK |
| 0 | Immune system | Adaptive_NK | TMEM244 | KLRC1,FCER1G,SYK |
| 0 | Immune system | Adaptive_NK | IER5L | KLRC1,FCER1G,SYK |
| 0 | Immune system | Adaptive_NK | RASGEF1A | KLRC1,FCER1G,SYK |
| 0 | Immune system | Adaptive_NK | MID2 | KLRC1,FCER1G,SYK |
| 0 | Immune system | Adaptive_NK | CD3D | KLRC1,FCER1G,SYK |
| 0 | Immune system | Adaptive_NK | SELM | KLRC1,FCER1G,SYK |
| 0 | Immune system | Adaptive_NK | TLR3 | KLRC1,FCER1G,SYK |
| 0 | Immune system | Adaptive_NK | RAB6B | KLRC1,FCER1G,SYK |
| 0 | Immune system | Adaptive_NK | LRRC75A | KLRC1,FCER1G,SYK |
| 0 | Immune system | Adaptive_NK | MIR4435-2HG | KLRC1,FCER1G,SYK |
| 0 | Immune system | Adaptive_NK | KLRC4 | KLRC1,FCER1G,SYK |
| 0 | Immune system | Adaptive_NK | LTBP4 | KLRC1,FCER1G,SYK |
| 0 | Immune system | Adaptive_NK | PERP | KLRC1,FCER1G,SYK |
| 0 | Immune system | Adaptive_NK | SCN8A | KLRC1,FCER1G,SYK |
| 0 | Immune system | Adaptive_NK | DUSP19 | KLRC1,FCER1G,SYK |
| 0 | Immune system | Adaptive_NK | EDNRB-AS1 | KLRC1,FCER1G,SYK |
| 0 | Immune system | Adaptive_NK | CRYBB3 | KLRC1,FCER1G,SYK |
| 0 | Immune system | Adaptive_NK | SCD5 | KLRC1,FCER1G,SYK |
| 0 | Immune system | Adaptive_NK | LOC102724094 | KLRC1,FCER1G,SYK |
| 0 | Immune system | Adaptive_NK | ELOVL4 | KLRC1,FCER1G,SYK |
| 0 | Immune system | Adaptive_NK | SPATA6L | KLRC1,FCER1G,SYK |
| 0 | Immune system | Adaptive_NK | TF | KLRC1,FCER1G,SYK |
| 0 | Immune system | Adaptive_NK | EFNA5 | KLRC1,FCER1G,SYK |
| 0 | Immune system | Adaptive_NK | SLC45A1 | KLRC1,FCER1G,SYK |
| 0 | Immune system | Adaptive_NK | CDKN2B-AS1 | KLRC1,FCER1G,SYK |
| 0 | Immune system | Adaptive_NK | OTOF | KLRC1,FCER1G,SYK |
| 0 | Immune system | Adaptive_NK | C1orf61 | KLRC1,FCER1G,SYK |
| 0 | Immune system | Adaptive_NK | CHRNE | KLRC1,FCER1G,SYK |
| 0 | Immune system | Adaptive_NK | TKTL1 | KLRC1,FCER1G,SYK |
| 0 | Immune system | Adaptive_NK | TSPAN2 | KLRC1,FCER1G,SYK |
| 0 | Immune system | Adaptive_NK | LOC101928988 | KLRC1,FCER1G,SYK |
| 0 | Immune system | Adaptive_NK | PARD6G | KLRC1,FCER1G,SYK |
| 0 | Immune system | Adaptive_NK | STXBP6 | KLRC1,FCER1G,SYK |
| 0 | Immune system | Adaptive_NK | TRPC3 | KLRC1,FCER1G,SYK |
| 0 | Immune system | Adaptive_NK | JAKMIP2 | KLRC1,FCER1G,SYK |
| 0 | Immune system | Adaptive_NK | HEY2 | KLRC1,FCER1G,SYK |
| 0 | Immune system | Adaptive_NK | TRIM46 | KLRC1,FCER1G,SYK |
| 0 | Immune system | Adaptive_NK | SNPH | KLRC1,FCER1G,SYK |
| 0 | Immune system | Adaptive_NK | HRASLS5 | KLRC1,FCER1G,SYK |
| 0 | Immune system | Adaptive_NK | SLC14A2 | KLRC1,FCER1G,SYK |

| | | | | |
|---|---|---|---|---|
| 0 | Immune system | Adaptive_NK | KRTAP5-AS1 | KLRC1,FCER1G,SYK |
| 0 | Immune system | Adaptive_NK | WWTR1 | KLRC1,FCER1G,SYK |
| 0 | Immune system | Adaptive_NK | SPTBN5 | KLRC1,FCER1G,SYK |
| 0 | Immune system | Adaptive_NK | KIR2DL2 | KLRC1,FCER1G,SYK |
| 0 | Immune system | Adaptive_NK | SLC14A1 | KLRC1,FCER1G,SYK |
| 0 | Immune system | Adaptive_NK | C16orf45 | KLRC1,FCER1G,SYK |
| 0 | Immune system | Adaptive_NK | TJP3 | KLRC1,FCER1G,SYK |
| 0 | Immune system | Adaptive_NK | ATP1A3 | KLRC1,FCER1G,SYK |
| 0 | Immune system | Adaptive_NK | ST8SIA1 | KLRC1,FCER1G,SYK |
| 0 | Immune system | Adaptive_NK | AGAP1 | KLRC1,FCER1G,SYK |
| 0 | Immune system | Adaptive_NK | KCCAT198 | KLRC1,FCER1G,SYK |
| 0 | Immune system | Adaptive_NK | PTPRM | KLRC1,FCER1G,SYK |
| 0 | Immune system | Adaptive_NK | NFIA | KLRC1,FCER1G,SYK |
| 0 | Immune system | Adaptive_NK | DGKH | KLRC1,FCER1G,SYK |
| 0 | Immune system | Adaptive_NK | GREB1 | KLRC1,FCER1G,SYK |
| 0 | Immune system | Adaptive_NK | LINC00565 | KLRC1,FCER1G,SYK |
| 0 | Immune system | Adaptive_NK | NUGGC | KLRC1,FCER1G,SYK |
| 0 | Immune system | Adaptive_NK | TUBB4A | KLRC1,FCER1G,SYK |
| 0 | Immune system | Adaptive_NK | IL5RA | KLRC1,FCER1G,SYK |
| 0 | Immune system | Adaptive_NK | SGCB | KLRC1,FCER1G,SYK |
| 0 | Immune system | Adaptive_NK | NPR3 | KLRC1,FCER1G,SYK |
| 0 | Immune system | Adaptive_NK | DENND2C | KLRC1,FCER1G,SYK |
| 0 | Immune system | Adaptive_NK | CD52 | KLRC1,FCER1G,SYK |
| 0 | Immune system | Adaptive_NK | CDC14C | KLRC1,FCER1G,SYK |
| 0 | Immune system | Adaptive_NK | MUC3A | KLRC1,FCER1G,SYK |
| 0 | Immune system | Adaptive_NK | ERRFI1 | KLRC1,FCER1G,SYK |
| 0 | Immune system | Adaptive_NK | LILRB1 | KLRC1,FCER1G,SYK |
| 0 | Immune system | Adaptive_NK | IMPG1 | KLRC1,FCER1G,SYK |
| 0 | Immune system | Adaptive_NK | GSC | KLRC1,FCER1G,SYK |
| 0 | Immune system | Adaptive_NK | SCUBE3 | KLRC1,FCER1G,SYK |
| 0 | Immune system | Adaptive_NK | WNT1 | KLRC1,FCER1G,SYK |
| 0 | Immune system | Adaptive_NK | EPHX2 | KLRC1,FCER1G,SYK |
| 0 | Immune system | Adaptive_NK | MAMLD1 | KLRC1,FCER1G,SYK |
| 0 | Immune system | Adaptive_NK | NACAD | KLRC1,FCER1G,SYK |
| 0 | Immune system | Adaptive_NK | LAMB1 | KLRC1,FCER1G,SYK |
| 0 | Immune system | Adaptive_NK | CHSY3 | KLRC1,FCER1G,SYK |
| 0 | Immune system | Adaptive_NK | SHB | KLRC1,FCER1G,SYK |
| 0 | Immune system | Adaptive_NK | ADAMTSL5 | KLRC1,FCER1G,SYK |
| 0 | Immune system | Adaptive_NK | LOC101929719 | KLRC1,FCER1G,SYK |
| 0 | Immune system | Adaptive_NK | TRPC1 | KLRC1,FCER1G,SYK |
| 0 | Immune system | Adaptive_NK | IL32 | KLRC1,FCER1G,SYK |
| 0 | Immune system | Adaptive_NK | ZFP36 | KLRC1,FCER1G,SYK |
| 0 | Immune system | Adaptive_NK | WDR74 | KLRC1,FCER1G,SYK |
| 0 | Immune system | Adaptive_NK | CD160 | KLRC1,FCER1G,SYK |
| 0 | Immune system | Adaptive_NK | XCL2 | KLRC1,FCER1G,SYK |
| 0 | Immune system | Adaptive_NK | ID2 | KLRC1,FCER1G,SYK |
| 0 | Immune system | Adaptive_NK | IER2 | KLRC1,FCER1G,SYK |
| 0 | Immune system | Adaptive_NK | CD7 | KLRC1,FCER1G,SYK |
| 0 | Immune system | Adaptive_NK | SNORD3A | KLRC1,FCER1G,SYK |
| 0 | Immune system | Adaptive_NK | CCL3 | KLRC1,FCER1G,SYK |
| 0 | Immune system | Adaptive_NK | TMEM107 | KLRC1,FCER1G,SYK |
| 0 | Immune system | Adaptive_NK | FOS | KLRC1,FCER1G,SYK |
| 0 | Immune system | Adaptive_NK | XCL1 | KLRC1,FCER1G,SYK |
| 0 | Immune system | Adaptive_NK | SNORD3D | KLRC1,FCER1G,SYK |
| 0 | Immune system | Adaptive_NK | MAP3K8 | KLRC1,FCER1G,SYK |
| 0 | Immune system | Adaptive_NK | JUN | KLRC1,FCER1G,SYK |
| 0 | Immune system | Adaptive_NK | PPP1R15A | KLRC1,FCER1G,SYK |
| 0 | Immune system | Adaptive_NK | VIM | KLRC1,FCER1G,SYK |

| | | | | |
|---|---|---|---|---|
| 0 | Immune system | Adaptive_NK | RNU12 | KLRC1,FCER1G,SYK |
| 0 | Immune system | Adaptive_NK | SNORD3B-1 | KLRC1,FCER1G,SYK |
| 0 | Immune system | Adaptive_NK | IL2RB | KLRC1,FCER1G,SYK |
| 0 | Immune system | Adaptive_NK | TMIGD2 | KLRC1,FCER1G,SYK |
| 0 | Immune system | Adaptive_NK | RP11-386I14.4 | KLRC1,FCER1G,SYK |
| 0 | Immune system | Adaptive_NK | GZMK | KLRC1,FCER1G,SYK |
| 0 | Immune system | Adaptive_NK | LTB | KLRC1,FCER1G,SYK |
